# Cell Painting and chemical structure read-across can complement each other for rat acute oral toxicity prediction in chemical early de-risking

**DOI:** 10.1101/2024.04.25.591123

**Authors:** Fabrice Camilleri, Joanna M. Wenda, Claire Pecoraro-Mercier, Jean-Paul Comet, David Rouquié

## Abstract

Early de-risking decisions in the development of new chemical compounds, enable the identification of novel chemical candidates with improved safety profiles. In vivo studies are traditionally conducted in the early assessment of acute oral toxicity of crop protection products to avoid compounds which are considered “very acutely toxic”, with an in vivo Lethal Dose of 50% (LD50) ≤ 60 mg/kg bodyweight. Those studies are lengthy, costly, and raise ethical concerns, catalyzing the use of non-animal alternatives. The objective of our analysis was to assess the predictive efficacy of read-across approaches for acute oral toxicity in rats, comparing the use of chemical structure information, in vitro biological data derived from the Cell Painting profiling assay on U2OS cells or the combination of both. Our findings indicate that the classification of compounds as very acute oral toxic (LD50 ≤ 60 mg/kg) or not is possible using a read-across approach, with chemical structure information, morphological profiles, or a combination of both. When classifying compounds structurally similar to those in the training set, chemical structure was more predictive (balanced accuracy of 0.82). Conversely, when the compounds to be classified were structurally different from those in the training set, the morphological profiles were more predictive (balanced accuracy of 0.72). Combining the two models allowed for the classification of compounds structurally similar to those in the training set, to slightly improve the predictions (balanced accuracy of 0.85).

For Table of Contents Only

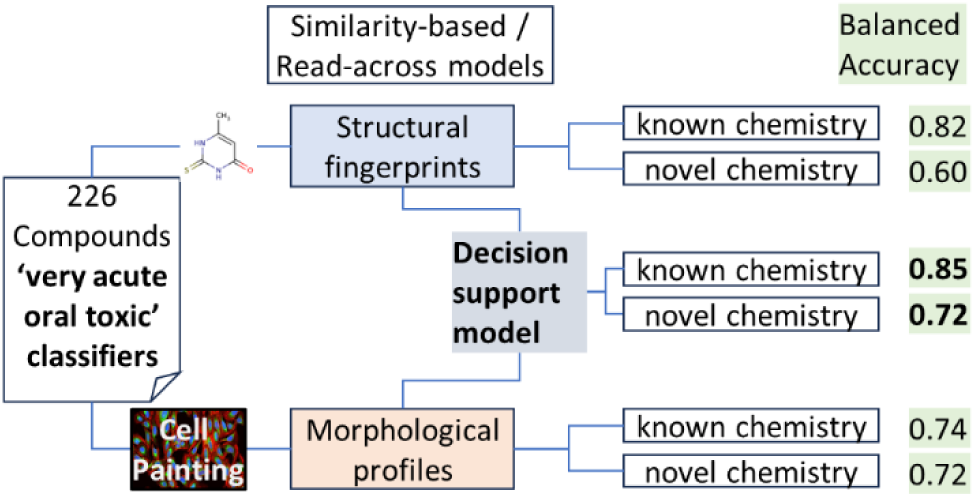

## Introduction

Small molecule discovery involves identifying and developing novel chemical compounds with optimized safety profiles across various target and non-target species, including laboratory animals, humans and environmental species. This complex process requires the integration of chemical and biological exploration. Chemists design diverse compounds to interact with specific biological targets or pathways while biological assays assess efficacy and safety, guiding further chemical optimization. Balancing potency with selectivity and minimizing off-target effects presents a significant challenge. To achieve this, chemists often explore new chemical spaces to discover novel structures with the desired properties^1,2^. Additionally, the complexity of biological systems poses hurdles, as the interplay of various factors influences a molecule’s properties. Success in small molecule discovery hinges on a multidisciplinary approach, where chemists, biologists and data scientists collaborate to navigate the intricate landscape of chemical and biological interactions, ultimately advancing the development of new active substances. In agrochemical discovery, early de-risking is crucial involving systematic assessment of compounds for potential safety issues to be addressed as early as possible. Addressing genotoxicity and acute oral toxicity is essential due to established cut-off criteria ^3^.

Traditionally, early genotoxicity evaluations are performed with in vitro test methods (Ames and Micronucleus assays) whereas acute oral toxicity profile screening involve laboratory animal testing in rodent to obtain a first estimate of the LD50 representing the single dose level at which lethality is induced in at least in 50 % of the tested animals. The traditional in vivo studies for acute oral toxicity assessment are time-consuming, expensive, and have low throughput, making it challenging to test a large number of chemicals. Animal studies also raise ethical concerns and must be reduced or if possible, eliminated. Higher throughput alternatives are desired, for the ranking of chemical candidates for their prioritization into the R&D pipeline based on their toxicological profiles. This minimizes resource waste, specifically extended R&D efforts for a chemical which is otherwise only later discovered to not meet required safety standards.

Non-animal alternatives are already available to reduce in vivo studies including in vitro approaches and in silico models for early estimation of the LD50. For instance, the in vitro 3T3 neutral red uptake assay (NRU) ^4^ can serve to categorize compounds with in vivo LD50 greater or lower than 2000 mg/kg. However, this LD50 threshold is high and may not discriminate compounds with lower LD50s. In acute toxicity testing, low LD50 value (LD50 ≤ 60mg/kg), means high potential for acute toxicity, which is an unwanted safety profile^3^.

In silico models are emerging which may close this gap. Quantitative structure activity relationship (QSAR) models rely on machine learning based on structural information of chemical compounds. One notable example is the Collaborative Acute Toxicity Modeling Suite (CATMoS), a public QSAR model, built in a collaboration of several research groups. CATMoS was trained on more than 10,000 compounds and demonstrated high performance in predicting acute oral toxicity in rats ^5^. As a regression model, CATMoS can predict the LD50 and classify chemicals into the five categories of Global Harmonized System (GHS). However, it is important to note that QSAR model predictions are only reliable within their applicability domain namely the chemical space covered by the chemistry represented in the training set. This represents a significant limitation when using such models to predict properties of novel chemistry not represented in the training set.

In this situation, we hypothesized that using in vitro high biologically dense profiles could be used instead of chemical structure information, especially when exploring new chemical spaces. The Carpenter–Singh Lab at the Broad institute has developed an such in vitro high content biologically profiling assay, Cell Painting, which capture the morphological information of cells perturbed by chemicals ^6^. The primary advantage of Cell Painting lies in its untargeted nature, theoretically allowing it to capture any bioactivity inducing a change in cell morphology. Additionally, this assay is more cost-effective than other profiling assays such as transcriptomics^7^. Cell Painting has already been used successfully, in ‘hit’ discovery and in Mode of Action (MoA) prediction ^8–10^. In the field of toxicology, the US Environmental Protection Agency (EPA) has explored its use to screen bioactive compounds for human risk assessment ^11^. More recently, Cell Painting has also been utilized for the prediction of mitochondrial toxicity^12,13^, and liver toxicity^14^.

To evaluate if biological information could complement compound structure predictions for acute oral toxicity especially when exploring new chemical spaces, classifiers were employed to predict whether compounds had very high acute oral toxicity (LD50 ≤ 60 mg/kg) or not (LD50 > 60 mg/kg). Initially, a well-performing public QSAR model, CATMoS, was utilized for the prediction of a set of Bayer Cros Science compounds. Secondly, a simple chemical structural similarity-based classifier (utilizing a K nearest neighbor ^15^) was employed for the prediction of the same set of compounds. This approach, also known as the read-across approach, is commonly used in toxicology specifically in the case of tox data poor chemicals, such as REACH compounds with limited number of in vivo results^16^. To simulate scenarios where predictions using the models are made on novel chemistry, a "novel chemistry" holdout strategy was created to assess the classifier. To verify the efficacy of this holdout strategy, the chemical structural similarity-based classifier was again evaluated. A Cell Painting assay was conducted on a smaller subset of compounds using the U2OS cell line to evaluate read-across using morphological profiles. The results of the two similarity-based classifiers were assessed: one based on chemical structure, and one based on morphological profiles derived from Cell Painting. Both classifiers were also tested in the context of the “novel chemistry” holdout strategy. Finally, a decision support model was constructed to determine which of the two similarity-based classifiers (chemical structure or morphological profile) should be recommended for prediction of acute toxicity.

Overall, our results showed that the classification of compounds as very acute oral toxic (LD50 ≤ 60 mg/kg) is possible using a read-across approach, with chemical structure information, morphological profiles, or a combination of both. When classifying compounds structurally similar to those of the training set, chemical structure was more predictive (balanced accuracy of 0.82). Conversely, when the compounds to be classified were structurally different from those of the training set, the morphological profiles were more predictive (balanced accuracy of 0.72). Combining both models allowed for the classification of compounds structurally similar to those used to train the classifiers, to slightly enhance the predictions (balanced accuracy of 0.85).

## Materials and methods

### Acute oral toxicity compound classes

The compounds were divided into two classes. The class designated as "Very acutely oral toxic" (abbreviated VAOT) included compounds with a lethal dose (LD50) of 60 mg/kg or less. The class designated as "Not very acutely oral toxic" (abbreviated NVAOT) included compounds with a lethal dose (LD50) greater than 60 mg/kg. (Table 1).

**Table 1.**
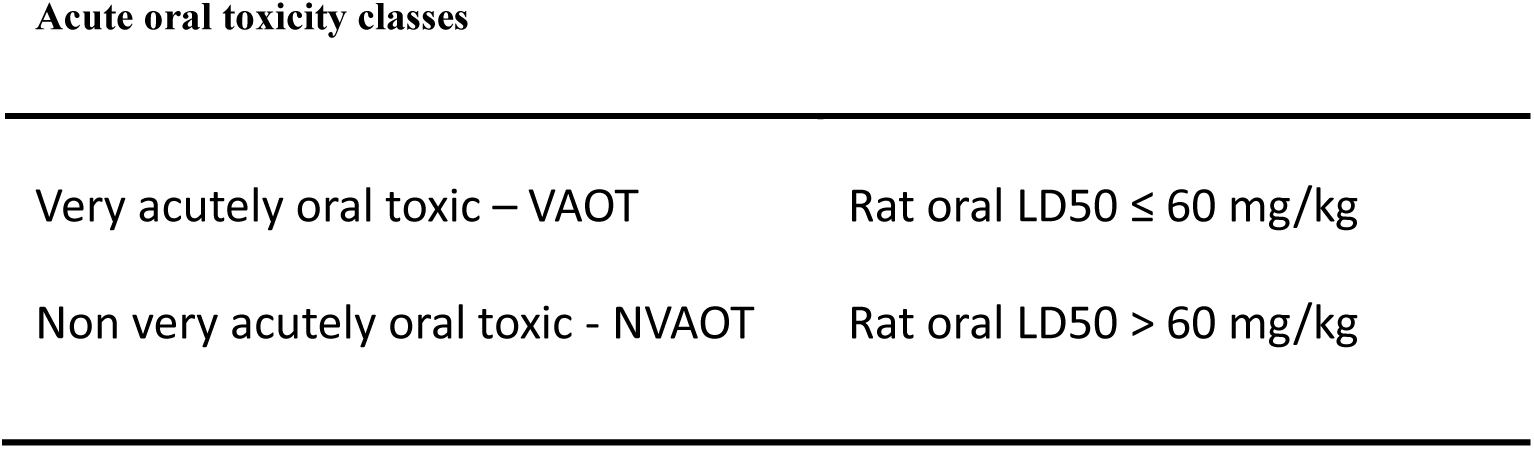
Definition of the two acute oral toxicity classes.

|  |  |
| --- | --- |
| Very acutely oral toxic – VAOT | Rat oral LD50 ≤ 60 mg/kg |
| Non very acutely oral toxic - NVAOT | Rat oral LD50 $>$ 60 mg/kg |

### Compound selection

#### Compounds with acute oral toxicity results in rats

To select compounds, Bayer internal databases were queried. A total of 765 compounds with in vivo rat acute oral toxicity results were found for two doses: 60 mg/kg and 300 mg/kg. Of the 765 compounds, 109 compounds were identified as acute toxic at the dose of 60 mg/kg, meaning compounds belonging to the VAOT class, and 521 compounds were not toxic at 300 mg/kg, meaning compounds belonging to the NVAOT class. To create a robust contrast between the morphological profiles of the VAOT and NVAOT classes, we excluded 135 compounds that were acute toxic at 300 mg/kg but not at 60 mg/kg. This resulted in the first dataset of 630 compounds, which were used to test the acute toxicity prediction of the collaborative Acute Toxicity Modeling Suite (CATMoS)^5^ and to build a chemical structure similarity-based classifier.

For the Cell Painting campaign, we checked which of the previous dataset compounds were available in Bayer internal compound repository, 81 VAOT compounds were found. To complete the list of VAOT compounds, we queried the chemIDplus public database ^17^, and selected 29 compounds that were available in Bayer compound logistics, making a total of 110 VAOT compounds. To have a balanced dataset, 116 NVAOT compounds were selected. To have a good chemical structure diversity among them, the Butina algorithm^18^ was used to cluster the 521 compounds of the previous dataset, based on the Tanimoto similarity of their Morgan fingerprint and a threshold of 0.7: a maximum of clusters was selected and the number of compounds coming from the same cluster were minimized (supplementary information Figure S1). This selection resulted in having a total of 116 NVAOT compounds and 110 VAOT.

To summarize, two sets of compounds were defined. The first one, called ‘QSAR only compound set’, was a set of 630 compounds; 109 were VAOT and 521 were NVAOT. The second one, called ‘Cell Painting compound set’, was a set of 226 compounds; 110 were VAOT, and 116 were NVAOT (Figure 1).

**Figure 1.**
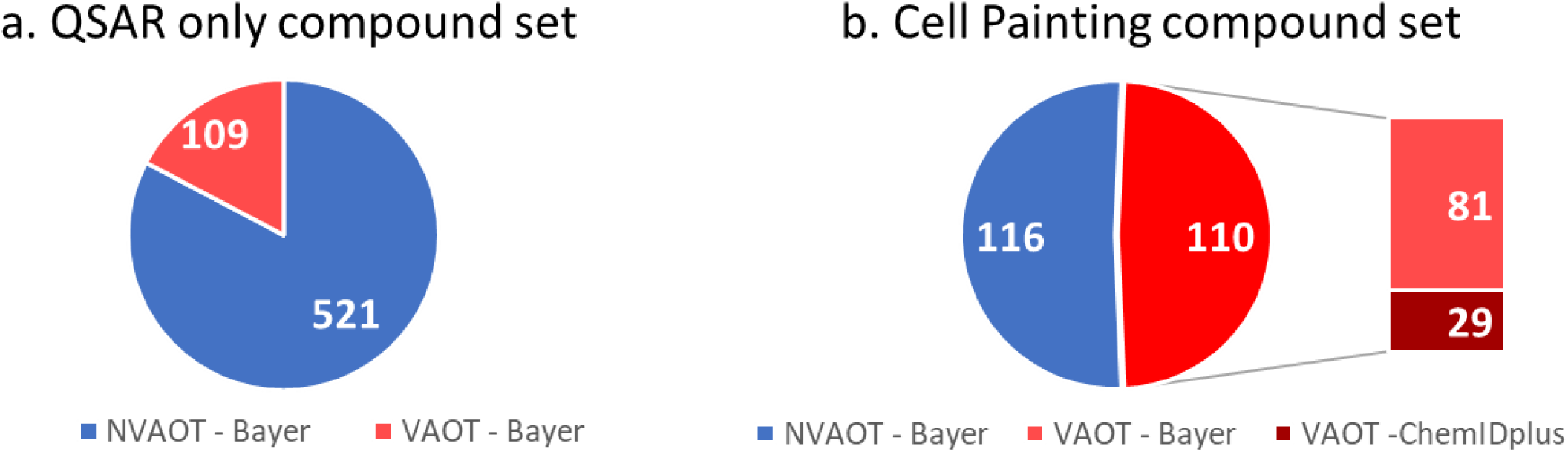
Composition of the two datasets by compound class: VAOT (Very Acutely Oral Toxic) and NVAOT (Non Very Acutely Oral Toxic); and by source: Bayer and ChemIDplus. a: QSAR only compound set. Set used for QSAR classifiers only. b: Cell Painting compound set, Set used for chemical structure and morphological profile classifiers.

#### Negative and positive controls

For the Cell Painting assay, DMSO (dimethyl sulfoxide) only (0.1%) was used as the negative control.

To monitor the Cell Painting assay’s performance and assess the quality of experiment’s replicates, a set of positive controls was used. These were compounds inducing reproducible and distinct morphological profiles in U2OS cells. The selection was based on published literature and pilot tests in our lab (Supplementary information).

### CATMoS QSAR model

As a first attempt, the acute oral toxicity class of the compounds was predicted using the collaborative Acute Toxicity Modeling Suite (CATMoS) QSAR model, implemented in the OPERA (version 2.9) ^5,19^ QSAR suite. To align with the two classes presented in this paper (VAOT and NVAOT), three CATMoS predictions were needed: the EPA classification, the GHS classification and the LD50 range estimation. All compounds being classified by CATMoS as EPA category 1 (LD50 ≤50 mg/kg), or GHS category 1 (LD50 ≤ 5 mg/kg), or GHS category 2 (5 mg/kg < LD50 ≤ 50 mg/kg) were designated as VAOT. For compounds being classified as EPA category 2 (50 mg/kg < LD50 ≤ 500 mg/kg), or GHS category 3 (50 mg/kg < LD50 ≤ 300 mg/kg), the lower limit of the LD50 range predictions was considered: if the inferior limits were smaller than 60, the compounds were classified as VAOT. For all other predictions, the compounds were classified as NVOAT.

The Opera implementation of CATMoS provides three prediction reliability metrics ^19^ that were used to understand the predictions made by the model. A global Applicability Domain Boolean value is calculated, indicating if a compound falls within the training set chemical space. Additionally, an Applicability Domain index is calculated, ranging from zero to one, revealing proximity of the queried compound to the training set. This index is relative to the similarity of the query chemical to its five nearest neighbors ^19^. In particular, a query chemical compound can belong to the CATMoS AD (global AD = 1), but can also be in a ‘gap’ of the training chemical space (AD index <0.6)^19^. In such cases, the predictions should be considered with caution^19^. Finally, a confidence index is computed, informing on the accuracy of the prediction of the neighbors of the queried compound.

### Cell painting campaign

A Cell Painting campaign was conducted in our laboratory to obtain the morphological profiles of our ‘Cell Painting compound set’ (set of 226 compounds). The Cell Painting Protocol v3 of the Broad Institute was utilized on U2OS human osteosarcoma cells with four biological replicates ^20^.

A previous Cell Painting pilot conducted at the unique dose of 10 µM demonstrated that few agrochemical compounds exhibited morphological changes compared to the negative control (see the ‘Morphological change signal measure’ section, and the ‘Biological response’ section). Therefore, in this campaign, to enhance the likelihood of capturing a morphological response, the compounds were screened at three concentrations: 10µM, 31.6µM and 100µM.

#### Cell Culture and Seeding

Human osteosarcoma cells U2OS have been purchased from ATCC (ref.: HTB-96, lot: 70025046). The McCoy’s 5A Modified Medium with GlutaMAX™ Supplement (Thermo Fisher, ref: 36600021) supplemented with 10% Fetal Bovine Serum (Gibco, ref.: 16000044) and penicillin/streptomycin mix (Sigma Aldrich, ref: P4458) was used for culturing cells in T75 or T175 flasks in a standard humidified incubator (37°C, 5% CO2). The passages were performed when the culture achieved about 80% confluency. Trypsin (Thermo Fisher, ref. 25200056) was used to detach the cells during passage and the number of live cells was calculated with an automatic cell counter (Countess II, Thermo Fisher) after staining the cells with trypan blue (Sigma, ref.: T8154). For the creation of a cell bank, the vial with frozen cells received from the supplier was thawed and expanded until internal passage no. 3 (P3). At this stage the cells were cryopreserved in complete media supplemented with 10% DMSO in an ultra-low temperature freezer (−150°C) creating a master bank. One vial of master bank was then thawed, expanded until internal passage no. 6 (P6), and cryopreserved as before to create a working bank. Vials of the working bank were then directly used for seeding the microplates. One vial of cells (containing 4 million cells) was removed from −150°C freezer and thawed in the water bath. The contents of the vial were immediately added to 10 ml of pre-heated complete media and centrifuged (5 min, 120xg). After removing the supernatant, the cell pellet was resuspended in 10 ml of complete medium through thorough pipetting. The cell suspension was then added to 150 ml of medium in a round bottle with a magnetic stirrer and immediately used for seeding the 384-well microplates (Greiner BioONE CELLSTAR µCLEAR®; ref: 781091). Multidrop (Thermo Fisher) was used to automatically distribute 36 µl of cell suspension per well, resulting in a seeding density of around 900 cells/well. The cells were then incubated at 37°C, in an atmosphere of 5% CO2 in an automatised incubator (Cytomat 2, Thermo Fisher). All experimental replicates were performed on a different day, using a separate cell vial originating from the same working bank (P6).

#### Chemical treatment

The test compounds were received in powder form in 96-well deep well plates. They were then dissolved in DMSO (dimethyl sulfoxide) to create 100 mM stock solutions, aliquoted in 96-well V-bottom plates (V96 PP Plate, Thermo Fisher) and frozen at −20°C until the day of the treatment. Every biological replicate of the experiment originates from a separate aliquot of the stock solution plate, so that the compounds undergo only 1 freeze-thaw cycle. On the day of the treatment (24 h post-seeding), the plates containing stock solutions were thawed and the compounds were diluted in DMSO to create dose plates containing 3 concentrations per compound: 100 mM, 31.6 mM, and 10 mM. The dilutions were performed with the use of Viper liquid handler (Synchron). The compound solutions form the dose plates were then administered to the cell plates in a two-step process. Firstly, an intermediate dilution was prepared: 1 µl of the compound solution was diluted in 100 µl of complete cell medium (1:100 dilution), next 4 µl of the resulting intermediate solution was administered to the cell plate (4 µl of the diluted compound into 36 µl of cell media, 1:10 dilution). The final concentrations of compounds that the cells were exposed to were therefore: 10 µM, 31.6 µM and 100 µM, the final vehicle (DMSO) concentration was 0.1%. The treated cell plates were subsequently incubated with the compounds for 48 h.

#### Staining

The staining and fixation were performed following the published protocol ^20^ with the use of PhenoVue JUMP kit (Perkin Elmer, ref.: PING23). Briefly, 20 µl/well of the Mitotracker solution were distributed to the cell plates with Multidrop (final concentration: 500 nM). After 30 minutes of incubation at 37°C, 20 µl/well of 16% PFA solution (Thermo Fisher, ref.: 28908) were added. The fixation was performed at room temperature (25°C). Two washes with HBSS buffer (Gibco, ref.: 14065-056) were performed with the aid of Mutlifo washer (BioTek). 20 µl/well of the staining solution (HBSS, 1% BSA, 0.1% Triton X-100, 43.7 nM PhenoVue Fluor 555 – WGA; 48 nM PhenoVue Fluor 488 - Concanavalin A; 8.25 nM PhenoVue Fluor 568 – Phalloidin; 1.62 µM PhenoVue Hoechst 33342 Nuclear Stain; 6 µM PhenoVue 512 Nucleic Acid Stain) were added and the plates were incubated for 30 min at room temperature before being washed again three times with HBSS. The plates were then sealed with aluminium foil and images were recorded directly.

### Morphological profile generation

#### Image acquisition

ImageXpress Micro 4 epifluorescent microscope (Molecular Devices) with 20x air objective was used for recording the fluorescent images (16-bit). The camera binning was set to 2×2. The total imaged area per well spanned 2163 µm x 2163 µm and consisted of 3 by 3 adjacent fields of view placed in the centre of the well. For each field of view images were recorded in 5 channels. The following filter sets were used: DAPI, GFP, Cy3, Texas Red, Cy5. The Z-offset and exposure times were set separately for each channel. A total of 207,360 images were acquired in this campaign.

#### Feature extraction

Morphological features were extracted using CellProfiler (version 4.2.1), the cell analysis software developed by the Broad Institute ^21^. Two CellProfiler pipelines were used: one pipeline for image illumination correction, and one pipeline for the image analysis. The image illumination correction works at the plate level and averages the intensity of the images of each channel. The image analysis pipeline, segmented objects on each image, labelled them according to the channel they were segmented on, and made thousands of measurements on those objects at the cell level. It also took measurement at the image level. A total of 4,761 features were measured and formed the morphological profile of a given cell.

#### Aggregation and normalization

After extracting the cell morphological profiles with CellProfiler, features were aggregated at the well level, by taking the means of each feature.

The features were then normalized, following the Broad Institute approach^6^, using the “mad robustized” method of the Python pycytominer package provided by the Broad Institute ^22^. The normalization process involved calculating the median of all wells on a plate for each feature, subtracting this value from the median absolute deviation (MAD) of the wells on the plate, and then multiplying the result by 1.4826 to obtain an unbiased estimator. To avoid a null denominator when the MAD was null, a value of 10^-18^ was added to the MAD.

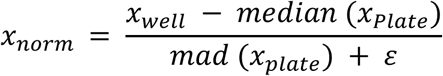

With: *x*_*norm*_ the normalized value of a morphological profile feature

*x*_*well*_ the value of a morphological profile feature

*x*_*Plate*_ the values of a morphological profile feature of a plate

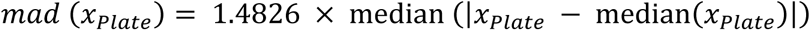

ε an infinitesimally small positive quantity (10^-18^) to avoid a null denominator.

### Quality check

To assess the quality of the Cell Painting experiment, several metrics were calculated. First, the number of cells of the negative control treatment (DMSO) was monitored, which was an output of the CellProfiler segmentation. The cell numbers should fall within the range of [1800; 3000] cells per well. The coefficient of variation of the number of cells for the negative control treatment should not exceed 15% per plate. All plates met the initial quality control standard.

To identify any other potential technical issues with the experiment, the Pearson correlations of positive controls across plates were calculated. The positive controls were selected to elicit very distinct and reproducible morphological profiles and were included in each plate. In a well-executed experiment, the replicates of these treatments should be well correlated. A correlation threshold of 0.8 was set for the Pearson correlation between replicates. Replicates were considered non well correlated if the correlation fell below this threshold. No outlier plates were identified at this stage, and all plates passed this quality control step.

There were 10 missing feature values at the well level in the morphological profiles. The majority of these values originated from the "Cells_AreaShape_FormFactor" feature. This feature was removed, along with two wells, in order to remove all missing values.

Further outliers were identified based on the number of cells within a group of replicates. Some wells exhibited a significant discrepancy in cell counts, with values exceeding 1800 cells, compared to other replicates of the same treatment. Twenty-two wells were identified as outliers and removed from the analysis.

### Unsupervised feature selection

To reduce the dimensionality of the normalized morphological profiles, an unsupervised feature selection was performed with the “feature_select” function of the pycytominer Python package ^22^.

This function performed several steps to select the features. First, highly correlated features were removed. For a pair of features with a Pearson correlation greater or equal to 0.9, the feature with the smallest sum of correlations with other features were removed. Second, features with low variances were removed. For a given feature, if the count of the second most common feature value divided by the count of the most common feature value was lower than 0.05, the feature was removed. Furthermore, features with the ratio defined as the number of unique feature values divided by the number of samples, below 0.01, were excluded. Third, a list of features (contained in the package), that are known to be noisy and generally unreliable were removed. Fourth, features with at least one absolute value greater than 500, values considered as outliers, were not retained. Finally, within each treatment group any features with a standard deviation greater than 1.2 were removed. This was done to identify and remove any noisy features. This process resulted in a total of 644 features, which were then used for downstream analysis.

### Consensus profiles

For a given treatment, in our case a chemical compound at a given concentration, the consensus profiles were obtained by aggregating the replicates. This was done by taking the median values of the replicates for each of the remaining features after the unsupervised feature selection.

### Morphological change signal measure

We utilized the Grit score^23,24^, a metric developed by the Broad Institute, to measure the morphological changes in a treatment replicate relative to the negative control treatment (DMSO). To calculate this metric, different Pearson correlation coefficients were calculated. First, the correlations between the morphological profiles of a given treatment replicate and each of the negative control treatments were calculated. The distribution of those correlations was defined by its mean and its standard deviation. Subsequently, the correlations between the morphological profiles of a given treatment replicate and other replicates of this treatment were calculated. Each of the previous correlation coefficients was z-transformed using the distribution of the correlations with the negative controls. The mean was subtracted, and the results were divided by the standard deviation. The grit scores were then obtained by taking the mean of the transformed values.

The grit score indicated how much a given replicate profile deviated from the negative control profiles. A high grit score indicated a high deviation of the profile from the negative control profiles.

Median grit scores for a given treatment were also calculated, taking the median of the treatment replicate grit scores. This median grit score value allowed measuring how much a treatment impacts the morphology of U2OS cells, compared to the negative controls (DMSO).

A threshold of 1 was set to indicate a treatment that induces a morphological change compared to the negative control. A grit score of 1 means that the correlation of the morphological profile of a treatment to its replicates, is one standard deviation away from the mean of its correlation with the negative control profiles.

### Molecular fingerprints

The compound structures were extracted from the Bayer database as SMILES format (Simplified Molecular-Input Line-Entry System). To reproduce the case when new chemical structures fall outside the applicability domain of chemical structure based predictive models the Morgan fingerprints were employed to describe the chemical structures of the compounds. To ensure sufficient Butina clusters for our data holdout strategy (see ‘Chemical compounds clustering with Butina’ and ‘Dataset splits section’), we selected a Morgan fingerprint on 1024 bits to provide the most detailed chemical structure description and thus differentiate more structural differences ^18,25,26^. To obtain them from the SMILES, we performed the following steps using the RDKIT Python package ^27^.

First, the SMILES were cleaned using the MolStandardize module of RDKIT. This involved removing hydrogens, disconnecting metal atoms, normalizing and ionizing the molecule, and keeping the parent fragments when several fragments of a compound existed. The molecule was then neutralized and, the canonical tautomer was returned. Finally, the cleaned SMILES were used to compute the Morgan fingerprints on 1024 bits, with a radius of three.

### Chemical compounds clustering with Butina

The Butina clustering algorithm groups molecules based on their structural similarity ^18^. The RDKit implementation of the Butina algorithm was used to cluster the chemical compounds ^27^. The clustering was based on the Tanimoto similarity of the Morgan fingerprints of the molecules, with a cut-off value of 0.7.

### Dataset splits

To assess the performance of the binary classifiers, the dataset was split several times into training and testing sets. Two types of splits were performed: a random one, which did not consider the chemical similarities of the compounds, and another split that aimed to create sets of structurally different chemicals. This was done to produce cases where the compounds to be classified were novel structures and therefore outside the Applicability Domain (Figure 2).

**Figure 2.**
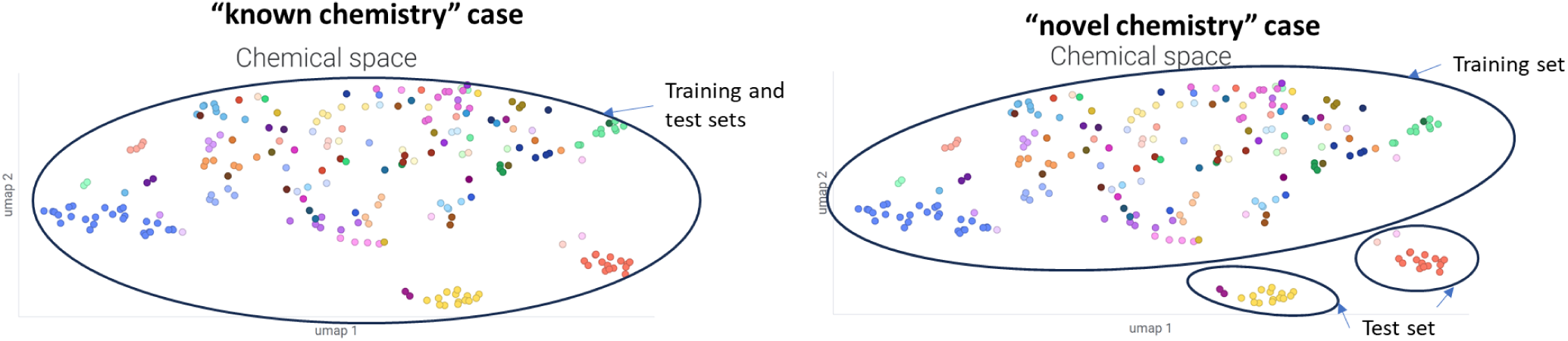
Example on an UMAP of the chemical space of training and test sets from the two holdout strategies. For the ‘known chemistry’ case, the training and test sets are originated from the same space. For the ‘novel chemistry’ case, the training and test sets are originated from different Butina clusters. Each color on the UMAP corresponds to a Butina cluster.

For the random split, called the ‘known chemistry case’, a stratified 10-fold-cross-validation was performed to split the dataset into 10 different training and testing sets. The scikit-learn Python package ^28^ was used to perform those splits with the StratifiedKFold function. The dataset was split 10 times with a 10-fold cross-validation with each cross-validation having a different random state, resulting in a total of 100 different splits. Each testing set included 22 or 23 compounds.

For the splits based on the chemical structures of the molecules, called the ‘novel chemistry case’, the compound structures were clustered using the Butina clustering algorithm ^18^. A cluster number was then assigned to each compound. The StratifiedGroupK-Fold function of scikit-learn was used to make a 10-fold cross-validation based on the cluster number ^28^. Indeed, this function assigned cluster numbers to the testing sets that differed from those in the training set. Additionally, it attempted to maintain a consistent ratio of VAOT and NVAOT classes within each set. The dataset was split in this manner 10 times, with different random states, resulting in a total of 100 unique splits. If any of these splits did not include compounds from both classes in the test sets, they were discarded.

### Binary classification classifier

To classify compounds as VAOT or NVOAT, several algorithms were tested. For this analysis, we decided to use a K Nearest Neighbors (KNN) algorithm ^15^, as it showed good performances (supplementary data). The KNN algorithm has also the advantage of being explainable and functions like a read-across, technique commonly used for toxicity prediction ^29^.

We used the scikit-learn ^28^ implementation of the K Nearest Neighbors (KNN) classification algorithm. Several classifiers were built depending on the data that were used as input. When using the chemical Morgan fingerprints, the Tanimoto (Jaccard) distance was used and when using the morphological profiles, the Pearson correlation-based similarity measure (1-Pearson correlation) was used. For all classifiers, we set the number of neighbors to one. The choice of the distances and the number of neighbors were the results of benchmarking done on both datasets (supplementary data).

### Decision support model

To aid the decision when the two types of classifiers (Morgan fingerprint and Cell Painting morphological profile classifiers) did not predict the same class, a model was built, similar to the Similarity-based merger model ^30^. This ensemble model takes as input the predictions of the two KNN classifiers, along with the distances of the nearest neighbors of each prediction (in total four values). A classifier was trained in each training set, for the cases where the two KNN classifiers did not agree on the predicted class. In the test sets, we used this model only when the two KNNs did not predict the same classes, otherwise the consensual predicted classes of the two classifiers were set as the final class (Figure 3).

**Figure 3.**
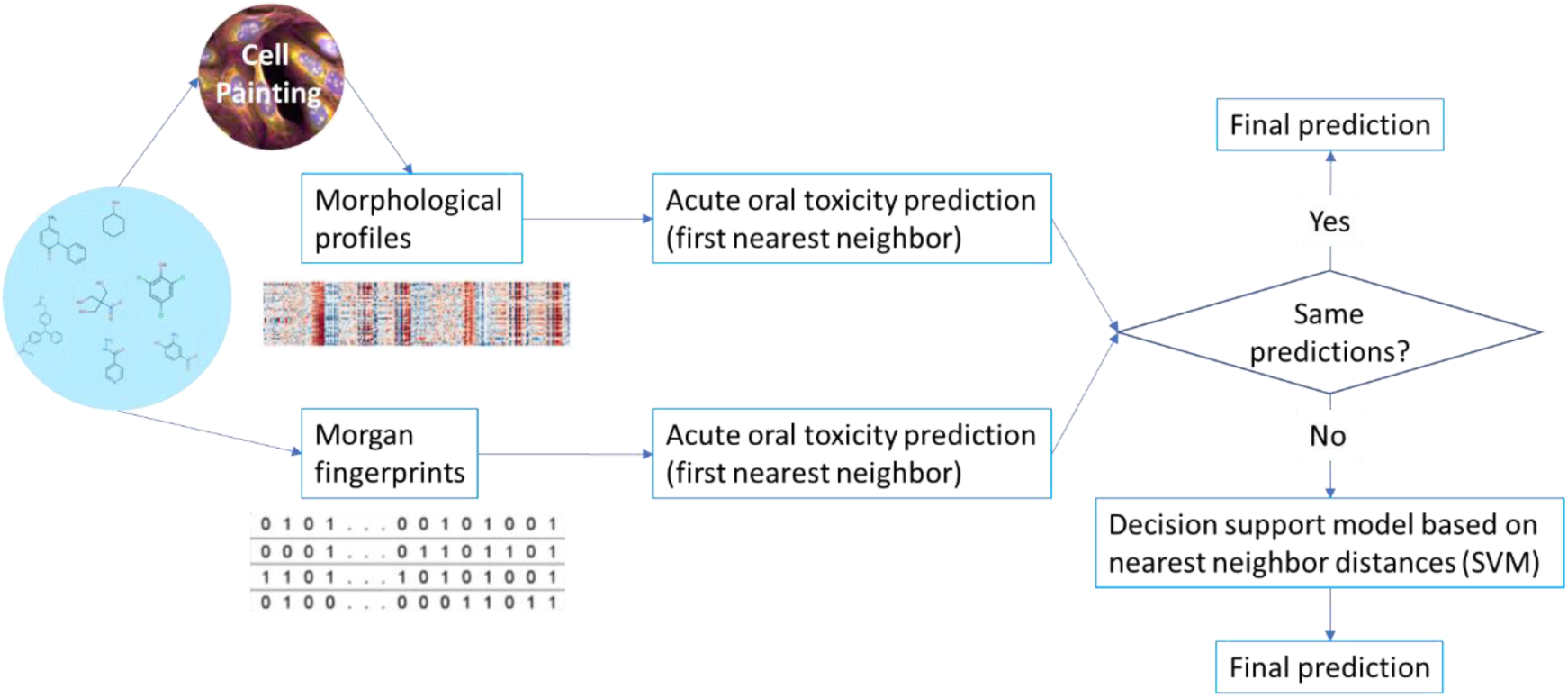
Decision support model

For this decision support model we used a SVM classifier ^31^ implementation of Scikit learn ^28^.

In the ‘novel chemistry’ case, the training set did not have enough examples with high distances. To remedy this, synthetic examples of distant structures were added. To do this, we subset the cases in each training set where the chemical structure-based predictions did not match the real class, and we updated the nearest neighbor distances with a random number between 0.7 and 0.9 and added these synthetic examples to the dataset used to train the model.

### Model performance evaluation

To evaluate the performance of the classifiers, we used different metrics. They were all based on the number of true positive (TP), true negative (TN), false positive (FP) and false negative (FN), which were the results of the model classification of a given testing set in a confusion matrix.

Sensitivity: 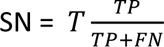

Specificity: 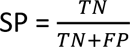

Balanced accuracy: 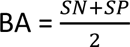

Matthews Correlation Coefficient: 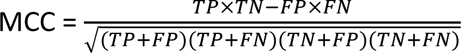

Accuracy: 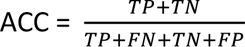

When using cross validation, those metrics were averaged over all testing sets, and their standard deviations calculated.

We performed a corrected t-test ^32^ to compare the balanced accuracy values over the splits of classification models.

### Visualization of the chemical and biological spaces

To visualize on two-dimensional scatter plots the chemical and the biological spaces, UMAP embeddings were generated using the Umap Python package ^33^. For the chemical space, the chemical compound structure similarities embeddings were calculated with the Tanimoto distances of the compound Morgan fingerprints. As for the biological space, the morphological profile similarities embeddings were computed using their Pearson correlation-based similarity measures. The plots were visualized in TIBCO Spotfire software.

## Results

The objective of our analysis was to assess the predictive efficacy of read-across approaches for acute oral toxicity in rats comparing the use of chemical structural information, in vitro biological data derived from the Cell Painting profiling assay on U2OS cells or the combination of both. Two distinct types of inputs were utilized to construct KNN models that classify compounds as Very acutely oral toxic (VAOT) or Non very acutely oral toxic (NVAOT).

Initially, we categorized 630 Bayer Crop Science (BCS) agrochemical compounds with known acute rat toxicity results using the public QSAR model CATMoS ^5^. Additionally, we used this specific unbalanced set of 630 compounds to build a structure similarity-based classifier.

Subsequently, KNN classifiers were used on a reduced but balanced set of 226 compounds using either the chemical structures or their morphological profiles in U2OS cells. A comprehensive analysis of both chemical space (represented by the chemical structures) and biological space (revealed by the U2OS morphological profiles) was conducted to enhance our comprehension of the classifier results. Finally, we investigated whether combining the predictions of the two classifiers could enhance the accuracy of the predictions.

The analysis demonstrated that a simple read-across approach, based on chemical structure information and biological data from the Cell Painting profiling assay on U2OS cells, can be used to predict acute oral toxicity, even in the context of new chemical space exploration.

### Results of the QSAR classifiers

Initially, CATMoS was employed to classify BCS compounds as either VAOT or NVAOT. We used the Opera CATMoS implementation of the model on the ‘QSAR only compounds set’ (Figure 1 1a), with 630 compounds as external test set. The CATMoS predictions were mapped to the two classes, with the majority of compounds being classified as NVAOT. Specifically, 5 of the 109 VAOT compounds, and 514 of the 521 NVAOT compounds were correctly predicted, resulting in a low sensitivity of 0.05, a high specificity of 0.99, a balanced accuracy of 0.52 and a MCC of 0.09 (Table 2a, supplementary data table S5). This outcome may be attributed to the fact that the CATMoS QSAR model was not trained on Bayer chemistry, but on mostly publicly available industrial chemical compounds. This indicates a possible mismatch in the applicability domain of the model for BCS chemistry.

**Table 2.**
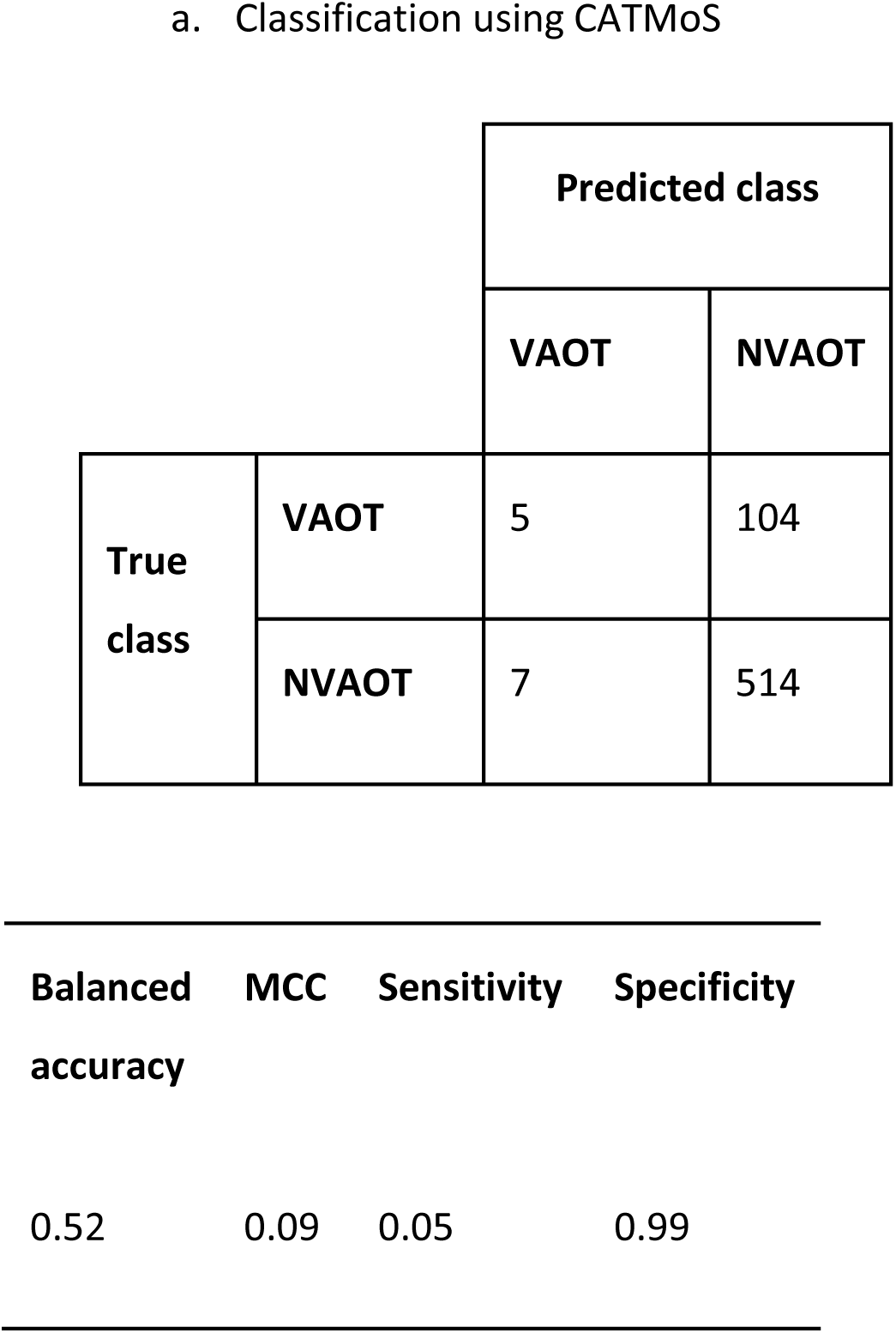
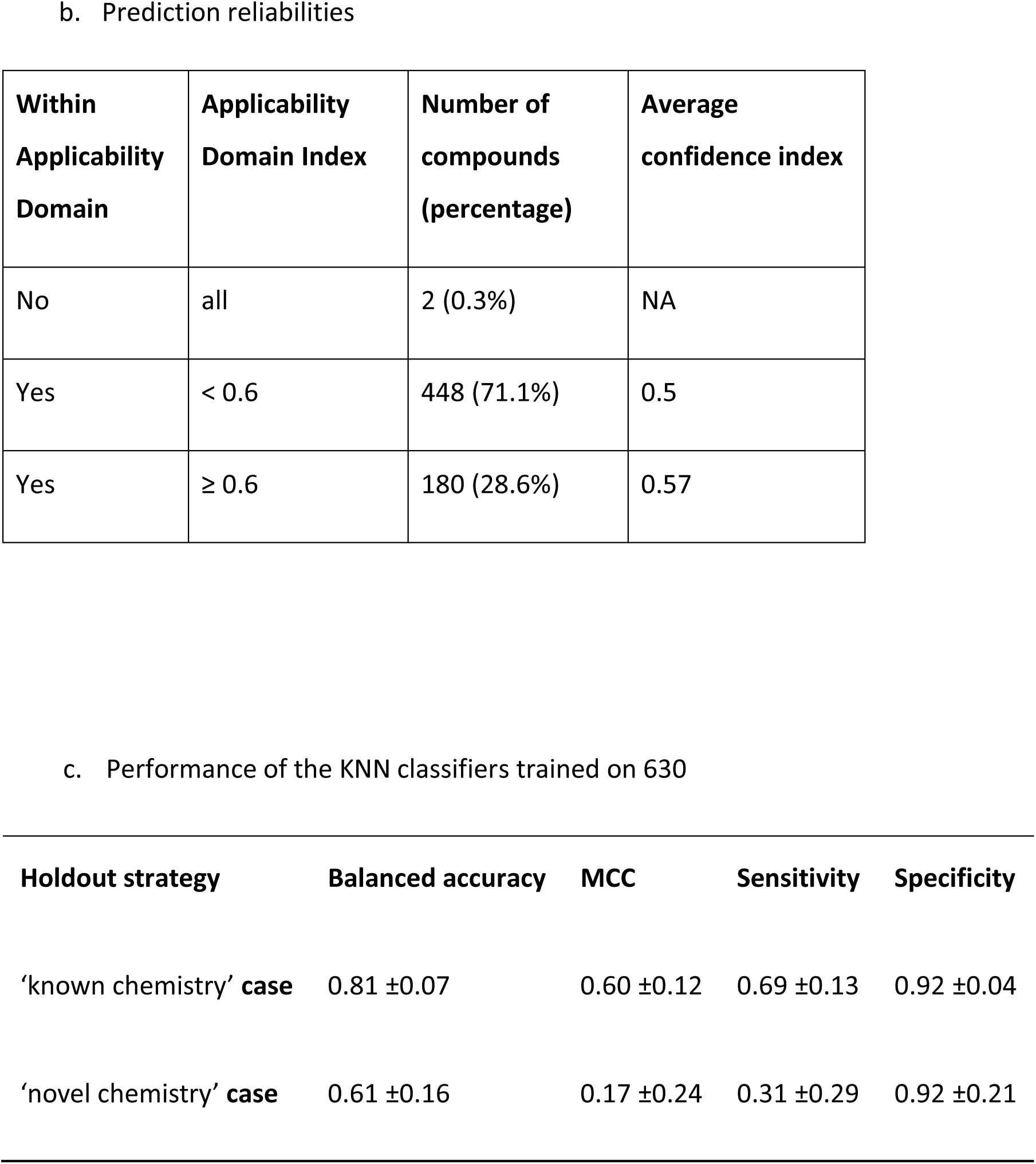
a. Confusion matrix and metrics for the classification of CATMoS for 630 compounds from Bayer Crop Science. b. Reliability of the predictions: Number of compounds outside the CATMoS applicability domain, number of compounds and average confidence index for compounds within the CATMoS applicability domain and having an Applicability Domain index below or above 0.6. c. Mean of 4 metrics assessing the performance of the KNN binary classifiers built out 630 Bayer CropScience agrochemical candidates, over the 100 splits of the ‘known chemistry’ case where training and testing sets are split randomly, not taking into account chemical structure similarities, and over the 44 valid splits of the ‘novel chemistry’ case where training and testing sets are split in order to have structurally different compounds over the two sets.

For the predictions of the set of 630 BCS compounds, we could determine that most compounds, 628 (99,6%) were within the CATMoS Applicability Domain (Table 2b). Most of them, 448 (71%), had an Applicability Domain index below 0.6, suggesting that the predictions should be considered with caution (Table 2c). The remaining 180 (29%) predictions having an Applicability Domain index above 0.6, displayed an average confidence level of 0.57, indicating a relatively low level of confidence in the predictions (Table 2b).

Subsequently, we developed a classifier, based on our 630 compound set, using a KNN classifier on Morgan fingerprints of chemical compounds. The classifier’s performance was evaluated through cross-validation with two data holdout strategies: the ‘known chemistry’ case, where compounds from the test sets resemble those from the training sets, and the ‘novel chemistry’ case, where compounds from the test sets differ structurally from those in the training sets.

In the ‘known chemistry’ case, the classifiers achieved an average balanced accuracy of 0.81, a sensitivity of 0.70, a specificity of 0.92 and a MCC of 0.61 (Table 2c).

In the ‘novel chemistry’ case, out of 100 theoretical cross-validation splits, only 44 of the 100 theoretical cross-validation splits included the two classes (VAOT and NVAOT) in both the training and testing sets, making them valid. The classifier demonstrated an average balanced accuracy of 0.60, a sensitivity of 0.33, a specificity of 0.87 and a MCC of 0.19 (Table 2c).

The ‘novel chemistry’ case demonstrated that chemical structure-based classifiers perform less well when classifying compounds that are structurally distinct from those in the training set. In summary, the chemical structure similarity-based models demonstrated good performance in handling known chemistry. However, as expected by the design of the ‘novel chemistry’ case, they exhibited a decrease in performance when confronted with unfamiliar chemical structures. This limitation becomes apparent when exploring new areas of chemical space. To address this limitation, we leveraged the biological effects of chemical compounds for predictions. The subsequent section outlines our approach, employing Cell Painting assay on U2OS cells to capture the biological effects of the chemical compounds.

### QSAR only compounds set (set of 630 compounds)

#### Comparison of chemical structure and Cell Painting morphological based classifiers

The objective of our study was to compare two inputs for predicting acute oral toxicity, utilizing a dataset called the ‘Cell Painting set’, a subset of the 630 ‘QSAR only compound set’, augmented with additional public chemical compounds. The Cell Painting set included a total of 226 compounds (Figure 1b). KNN classifiers were trained using both types of input and employing two data holdout strategies: ‘known chemistry’ and ‘novel chemistry’ cases.

Similarly to the previous chemical structure similarity-based classifier on the QSAR only compound set (630 compounds), KNN classifiers were trained on the Morgan fingerprint of the molecules.

For classifiers based on the morphological profiles obtained from Cell Painting, consensus profiles were utilized after normalization, unsupervised features selection, and replicate profile aggregation at the treatment level. Regarding the chemical structure similarity-based classifiers, we used KNN algorithm. Classifiers were built for each tested concentration (10 µM, 31.6 µM, 100 µM).

#### Results in the ‘known chemistry’ Case

In the ‘known chemistry’ case, the chemical structure similarity-based classifier demonstrated superior performance compared to other classifiers, achieving a mean balanced accuracy of 0.82. This was followed by the 31.6µM morphological profile classifier, with a mean balanced accuracy of 0.74 (Table 3a). The two other morphological profile classifiers at 10 and 100 µM demonstrated lower performance. (Table 3a).

**Table 3.** Mean and standard deviations of 4 metrics: Balanced Accuracy (BA), Matthew’s correlation coefficient (MCC), Sensitivity (SN), Specificity ‘SP). Four different input data were used to classify compounds as VAOT and NVAOT: chemical structural data (Morgan FP) and Cell Painting (CP) morphological profiles of U2OS cells exposed to chemicals at 3 concentrations (MP 10 µM, MP 31.6 µM and MP 100 µM). Orange highlight = the best average metric. The decision support (DS) model performance (combining the Morgan Fingerprint and the morphological profile 31.6 µM classifier predictions, supplemented with synthetic examples), are shown in the last row. Section a. Reports the performance of the binary classifiers, over the 100 splits of the ‘known chemistry’ case where training and testing sets are split randomly, not considering chemical structure similarities. Section b.Reports the performance of the KNN classifiers, over the 99 valid splits of the ‘novel chemistry’ case where training and testing sets are split to have structurally different compounds over the two sets.

|  | a: known chemistry case |  |  |  | b: novel chemistry case |  |  |  |
| --- | --- | --- | --- | --- | --- | --- | --- | --- |
|  | BA | MCC | SN | SP | BA | MCC | SN | SP |
| **Morgan FP** | 0.82 ± 0.08 | 0.65 ± 0.16 | 0.82 ± 0.11 | 0.82 ± 0.12 | 0.60 ± 0.16 | 0.20 ± 0.29 | 0.49 ± 0.28 | 0.71 ± 0.18 |
| **CP 10 μM** | 0.57 ± 0.08 | 0.14 ± 0.16 | 0.55 ± 0.14 | 0.58 ± 0.14 | 0.50 ± 0.14 | 0.04 ± 0.26 | 0.46 ± 0.24 | 0.54 ± 0.15 |
| **CP 31.6 μM** | 0.74 ± 0.08 | 0.49 ± 0.17 | 0.70 ± 0.13 | 0.78 ± 0.12 | 0.72 ± 0.12 | 0.42 ± 0.23 | 0.66 ± 0.22 | 0.78 ± 0.15 |
| **CP 100 μM** | 0.63 ± 0.09 | 0.27 ± 0.19 | 0.61 ± 0.13 | 0.65 ± 0.14 | 0.53 ± 0.13 | 0.07 ± 0.25 | 0.47 ± 0.23 | 0.60 ± 0.18 |
| **DS** | 0.85 ± 0.07 | 0.71 ± 0.14 | 0.86 ± 0.11 | 0.84 ± 0.11 | 0.72 ± 0.13 | 0.42 ± 0.26 | 0.67 ± 0.23 | 0.77 ± 0.16 |

The distribution of the balance accuracy values over the 100 splits for each input type of input showed a narrow range (Figure 4a), which was confirmed by low standard deviations ranging from 0.08 to 0.09 (Table 3a).

**Figure 4.**
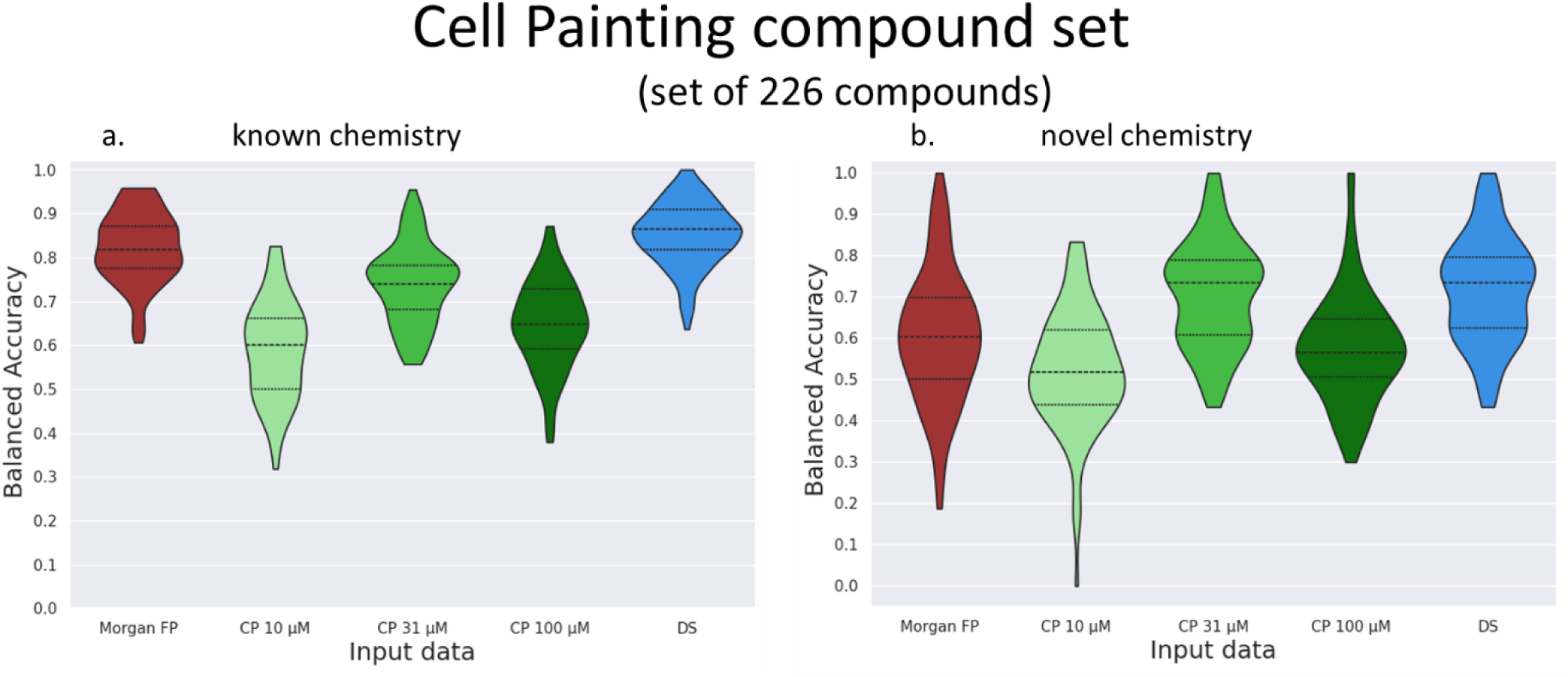
Section a: violin plots represent the balanced accuracies of the binary classifier for the 10 x 10-fold cross validation splits not considering the structure similarities (known chemical case). Section b: violin plots represent the balanced accuracies of the KNN binary classifier for 99 valid splits of the 10 x 10-fold cross validation that put in the testing set chemical structurally different from the training set (novel chemical case). Legend: In red, the classifier using the Morgan Fingerprint, in light green the classifier using the morphological profiles at 10 µM, in mid- green the classifier using the morphological profiles at 31.6 µM and in dark green, the classifier using the morphological profiles at 100 µM, in blue, the decision support (DS) model. Inside each violin plot the quartiles are indicated as dash lines.

A statistical analysis using the Nadeau and Bengio’s corrected t-test to compare the balance accuracy values over the 100 splits of the two top classifiers, indicated that the chemical structure similarity-based classifier significantly outperformed the 31.6 µM morphological profile classifier (p-value= 0.05).

#### Results in the ‘novel chemistry’ Case

In the ‘novel chemistry’ case, the 31.6 µM morphological profile classifier demonstrated superior performance achieving a mean balanced accuracy of 0.72. This was followed by the chemical structure similarity-based classifier, with a mean balanced accuracy of 0.60 (Table 3b). The remaining two classifiers demonstrated lower performance (Table 3b).

The distributions of the balanced accuracies for each classifier showed that in certain splits, the chemical structure similarity-based classifier encountered challenges in making accurate predictions (Figure 4b). This was also the case, to a lesser extent for the 31.6 µM morphological profile-based classifiers (Figure 4b).

The Nadeau and Bengio’s corrected t-test indicated that the 31.6 µM morphological profile classifier significantly outperformed the chemical structure similarity-based classifier (p- value=0.045).

In summary, for the chemical structure similarity-based classifiers, we reproduced the results of the previous chemical structure similarity based classifier, which was trained on approximately three times more compounds (630 compounds) with good performances in the ‘known chemistry’ case, and a decrease in performance in the ‘novel chemistry’ case (the balanced accuracy dropped from 0.82 to 0.60).

Overall, our findings emphasize the superior performance of chemical structure similarity-based classifier in the ‘known chemistry’ case. However, the morphological profile-based classifier remains a valuable tool, particularly in the ‘novel chemistry’ case, where the classifiers based on the 31.6 µM morphological profile demonstrated the highest performance (balanced accuracy of 0.72).

### Performances of the classifiers

#### Comparison of the chemical and biological spaces

To gain insight into the performance of the classifiers based on the chemical structures and the morphological profiles, we investigated into the chemical and biological spaces, for the Cell Painting set of compounds.

#### Chemical space

Our dataset comprises 226 compounds primarily originating from Bayer Crop Science chemistry and supplemented with 29 public compounds. The Butina algorithm, using the Tanimoto distance and a threshold of 0.7, identified 91 clusters. Of these, 61 were represented by a unique compound indicating structural diversity.

By plotting the similarity of the structures on a UMAP (Figure 5), using Morgan fingerprints and Tanimoto distance, it is possible to visually identify distinct clusters. Notably, certain clusters exclusively comprised of VAOT compounds (e.g., cluster A), while others were exclusively comprised of NVAOT compounds (e.g., cluster B). Additionally, several clusters contained a mixture of both (e.g., clusters C and D).

**Figure 5.**
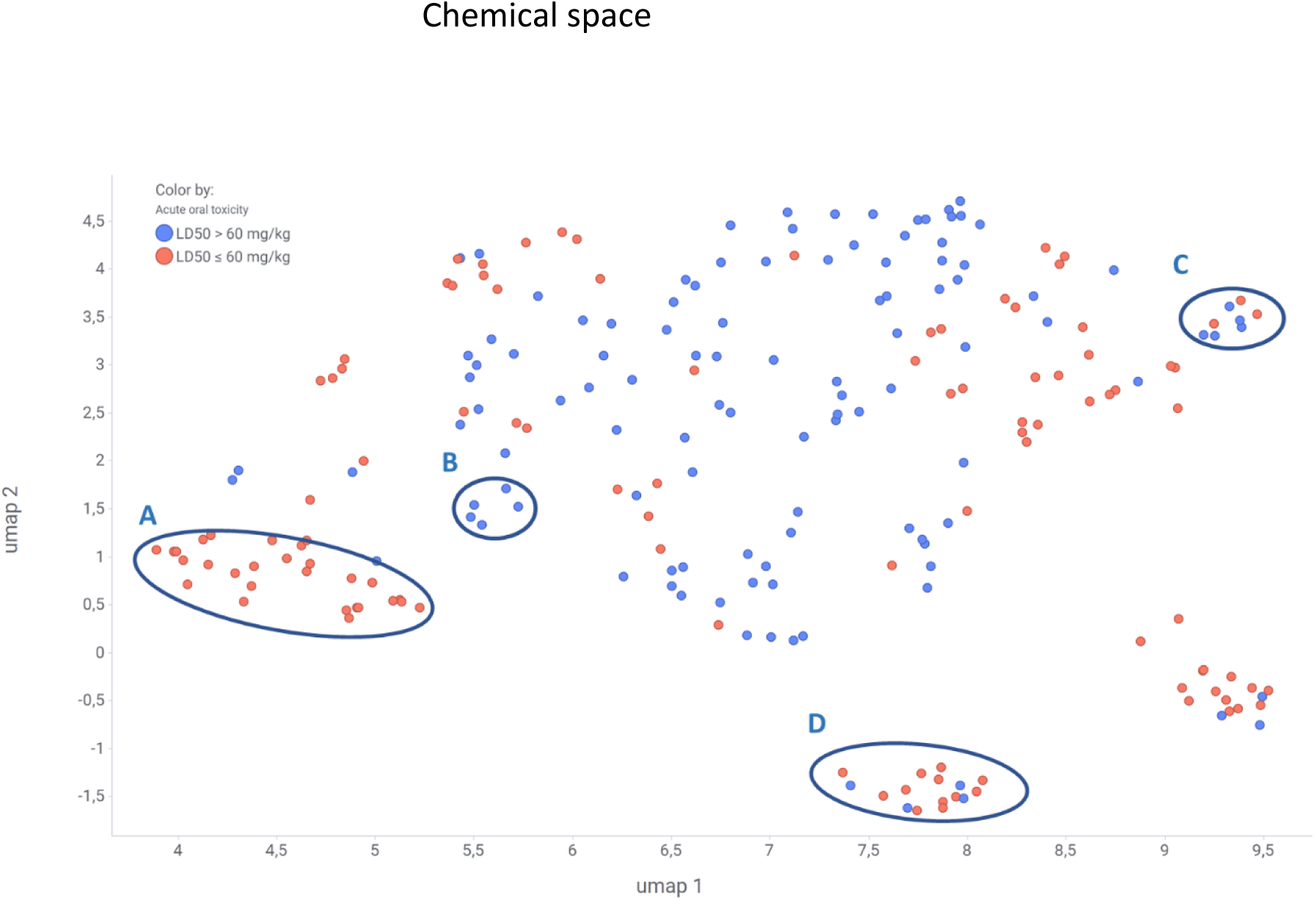
Scatter plot of the 2-dimensional UMAP embedding of the chemical compound morgan fingerprints. In blue, the chemical compounds that are NVAOT. In red, chemical compounds that are VAOT. Four clusters of compounds are designated by the letter A, B, C and D. Cluster A is an example of a cluster with only VAOT compound. The cluster B is an example of a cluster with only NVAOT compounds. B and C are two examples of clusters with a mix of VAOT and NVAOT compounds.

In summary, the chemical space exhibited diversity in the form of different clusters. Specific areas demonstrated a prevalence of either VAOT or NVAOT compounds, while others presented a combination of both classes.

#### Biological space

Figure 6 illustrates the degree of similarity in the biological response of compounds, as measured by Pearson correlation-based similarities, on a UMAP plot. In contrast to the chemical space, a limited number of clusters visually emerged with only two notable clusters observed. An isolated small cluster (cluster A) was clearly separated from the other profiles, and upon inspection these profiles corresponded to instances with notably low cell count.

**Figure 6.**
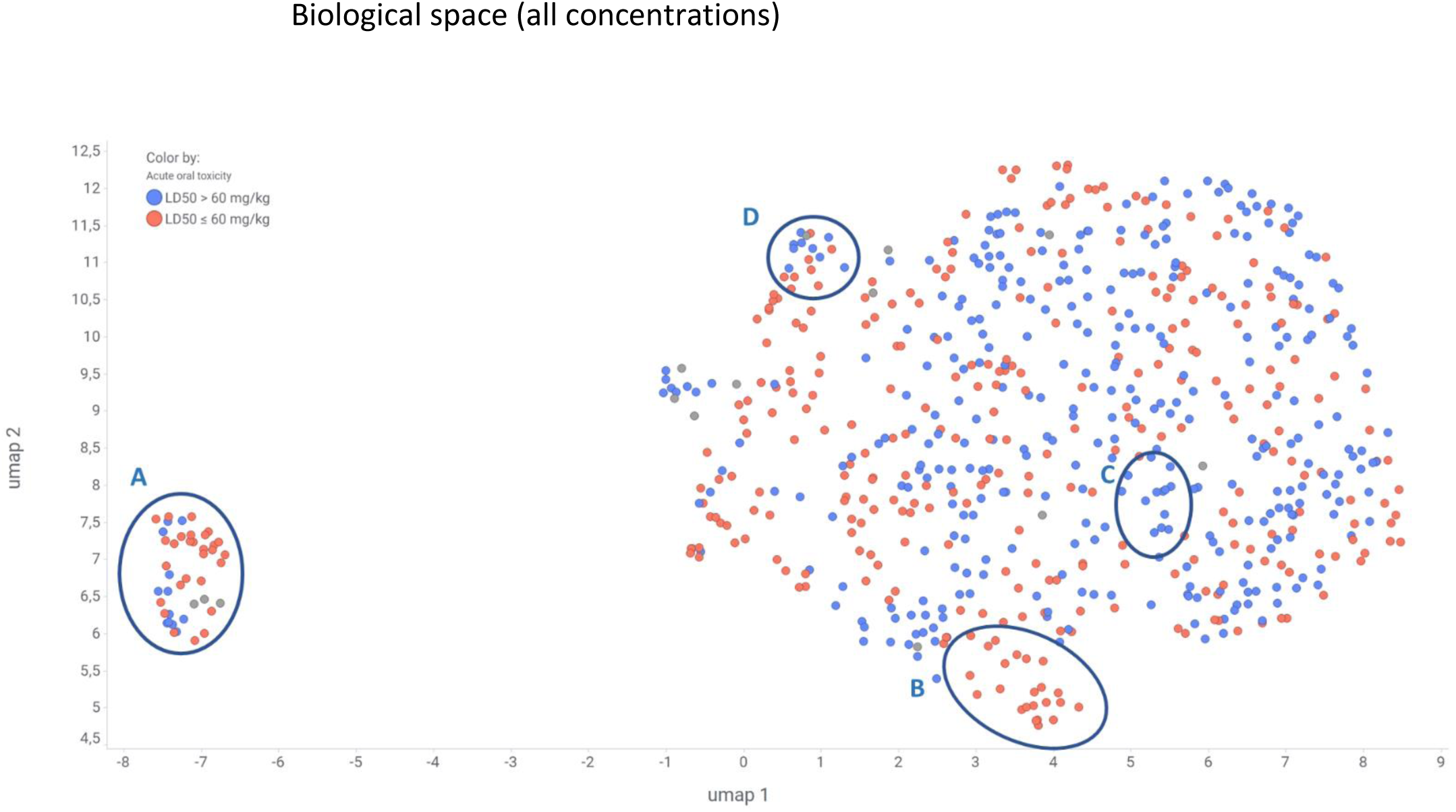
Two-dimensional representation of the consensus morphological profile similarities for all treatments and 3 concentrations, using Uniform Manifold Approximation (UMAP) embedding on 2 components with the Pearson correlation-based similarity measure. In red, the morphological profiles of U2OS cells perturbated by VAOT compounds. In blue, the morphological profiles of U2OS cells perturbated by NVAOT compounds. Four groups of compounds are designated by the letter A, B, C and D. The group A of compounds corresponds to treatment with very low cell counts. The group B of compounds is an example of grouping with a high number of VAOT compounds. The group C of compounds is an example of grouping with a high number of NVAOT compounds. The group D of compounds is an example of grouping with a mix of VAOT and NVAOT compounds.

The second cluster exhibited diverse areas: including regions with a high number of VAOT (e.g., grouping B), NVAOT compounds (e.g., grouping C), and areas with a mix of both classes (e.g., grouping D).

In Figure 7, we focused on the 31.6µM concentration, the concentration, which yielded optimal performance for the classifier using morphological profiles. Similar observations were made, with an isolated cluster corresponding to profiles with a very low number of cells. Additional distinct areas emerged, showcasing regions with high number of VAOT or NVAOT compounds, as well as areas with a mixed representation of both classes.

**Figure 7.**
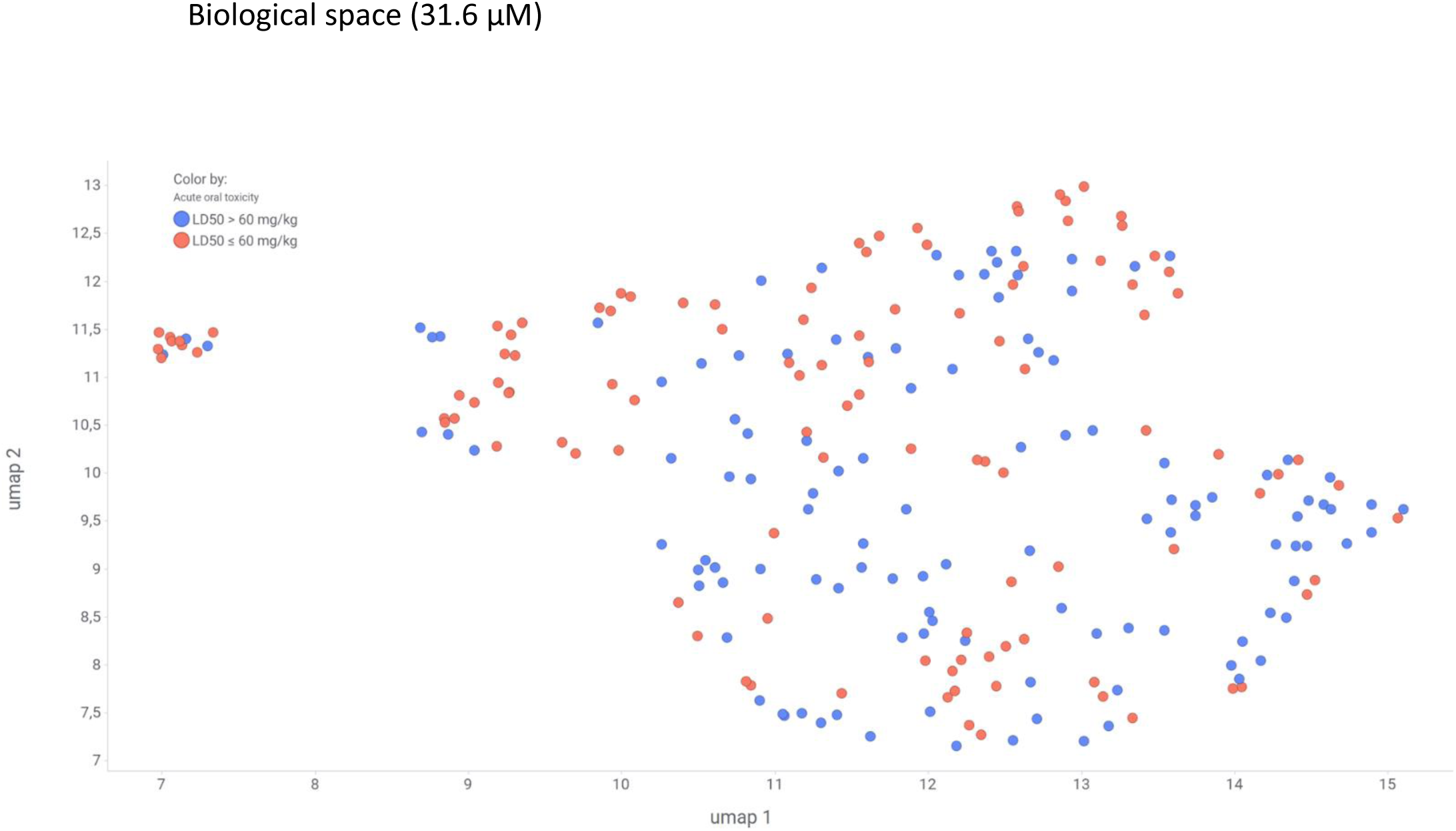
Two-dimensional representation of the consensus morphological profile similarities for all treatments at 31.6 µM, using Uniform Manifold Approximation (UMAP) embedding on 2 components with the Pearson correlation-based similarity measure. In red, the morphological profiles of U2OS cells perturbated by VAOT compounds. In blue, the morphological profiles of U2OS cells perturbated by NVAOT compounds.

#### Comparison of the chemical and biological spaces

We compared different groups of compound structures and groups of morphological profiles to better understand the chemical and biological space interrelationship. Notably, we observed that chemicals clustering together in the chemical space could elicit a diversity of biological responses in the biological space, emphasizing that structurally similar compounds may manifest distinct biological responses (Figure 8, left column). We illustrate a specific case involving a group of chemicals, the carbamates, inducing similar morphologies in U2OS cells (Figure 8, center column). Interestingly, similar morphological profiles induced by structurally different compounds could also be observed (Figure 8, right column). This comparison illustrates that: 1) the biological effects of structurally similar molecules may not necessarily be identical, 2) Structurally similar compounds could trigger different biological effects and 3) conversely, compounds with different structures could result in comparable biological responses.

**Figure 8.**
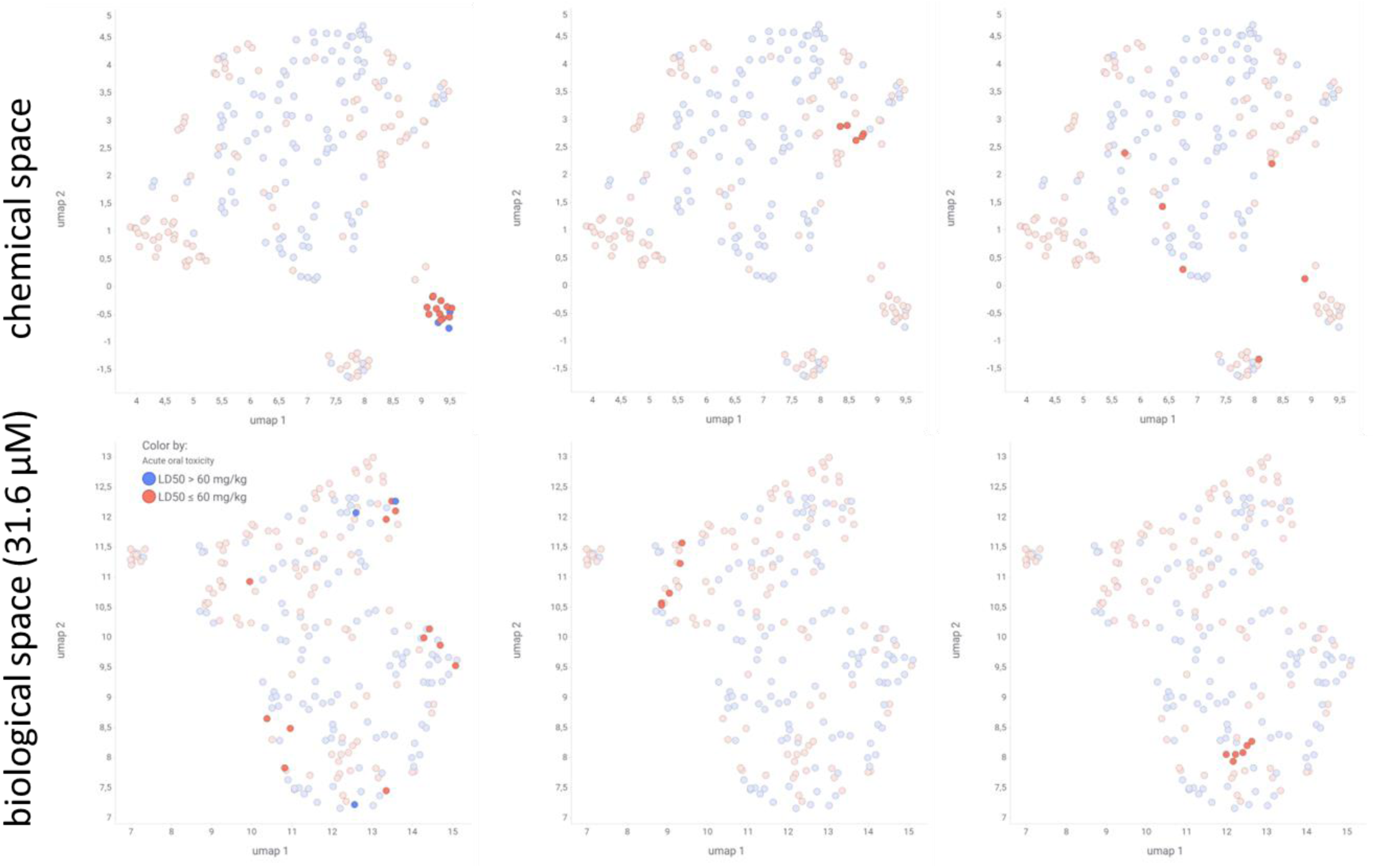
(top) Chemical space. Scatter plot of the 2-dimensional UMAP embedding of the chemical compound morgan fingerprints. (bottom) Biological space. Scatter plot of the 2-dimensional UMAP embedding of the Cell Painting morphological profiles of the chemical compounds at 31.6 µM. In red, the morphological profiles of U2OS cells perturbated by VAOT compounds. In blue, the morphological profiles of U2OS cells perturbated by NVAOT compounds. On each column, the chemical profiles and morphological profiles of the same compounds are selected. (left) Example of structurally similar compounds, inducing different morphological profiles. (center) Example of structurally similar compounds (carbamates), inducing similar morphological profiles. (right) Example of structurally different compounds, inducing similar morphological profiles.

#### Biological response

To analyze the biological response of U2OS cells to chemical compound perturbations, we employed morphological profiles, using two metrics: the Grit score ^22^, and the number of cells. The analysis was conducted to better understand how U2OS cells reacted to our set of compounds, which in turn helped us to understand the results of the classifiers.

#### Grit score

The Grit score indicated the extent to which the average morphology of U2OS cells perturbed by a compound deviated from the average negative control morphology of non-perturbed U2OS cells. A high Grit score indicated a more distinct cell morphology from the negative controls. For example, the average Grit score of the positive controls was 4.8.

A Mann-Whitney U rank test on the grit scores, for the two compound groups, VAOT and NVAOT, demonstrated that VAOT compounds elicited a marginal though significant stronger biological response compared to NVAOT (Grit score respectively 3 and 2.5, p-value of 0.004). (Table 4).

**Table 4.** Average grit score and number of cytotoxic treatments for different groups of profiles: VAOT (very acutely oral toxic compounds), NVAOT (non-very acutely oral toxic compounds), 10 µM treatment profiles, 31.6 µM treatment profiles and 100 µM treatment profiles.

| **profiles** | **Average**<br>**Grit score** | **Number of cytotoxic**<br>**treatments** |
| --- | --- | --- |
| **Negative control** | NA | 0 |
| **VAOT** | 3 | 28 |
| **NVAOT** | 2.5 | 16 |
| **10 µM all** | 1.9 | 11 |
| **10µM - VAOT** | 2.1 | 8 |
| **10µM - NVAOT** | 1.64 | 3 |
| **31.6 µM all** | 2.6 | 22 |
| **31.6µM - VAOT** | 2.98 | 14 |
| **31.6µM - NVAOT** | 2.17 | 8 |
| **100 µM all** | 3.7 | 44 |
| **100µM-VAOT** | 4.01 | 27 |
| **100µM-NVAOT** | 3.20 | 17 |

Regarding concentrations, on average, the 10 µM treatments had a grit score of 1.9, the 31.6 µM had a grit score of 2.6 and the 100 µM treatment had a grit score of 3.7. This aligns with our assumption that higher concentrations lead to increased biological responses, a consideration made when designing the Cell Painting campaign with 3 concentrations (Table 4).

Identifying compounds with no induced morphological changes, we set a Grit score threshold of 1. Below this threshold, we considered a treatment not inducing any morphological change. Of the 23 compounds falling below this threshold 6 were VAOT.

#### Number of cells

An additional output of the image analysis was the number of cells per well. For this analysis the number of cells was not normalized, and the median number of cells per well for a given treatment were computed. The average number of cells for the negative controls was 2231. We arbitrarily set the number of cells that defines cytotoxicity as a cell count below 50% of the average negative control cell count, meaning a cell count per well of below 1115 defined a cytotoxic treatment.

In total 44 compounds exhibited cytotoxicity: 11 compounds, at 10 µM, 22 compounds, at 31.6 µM and 44 compounds, at 100 µM (Table 4).

Categorizing by class, 28 VAOT compounds (25%) and 16 NVAOT compounds (14%) displayed cytotoxicity for at least one concentration. A chi-square test of independence of variables, with the null hypothesis that the number of cytotoxic compounds is independent of the class (VAOT and NVAOT) gave a p-value of 0.1. We could not conclude that there was a higher percentage of cytotoxicity for VAOT compounds (Table 4).

#### Results of the decision-support model

The decision-support model aided the decision when the two KNN classifiers did not predict the same class. The model combined four pieces of information: predictions from the KNN classifier based on the chemical structure information, predictions from the KNN classifier based on the morphological profiles and distances to the nearest neighbor in each classifier.

In the ‘known chemistry’ case, the model demonstrated an average balanced accuracy of 0.84, slightly above the chemical structure similarity-based classifier’s average balanced accuracy of 0.82 but was not significantly different (Nadeau and Bengio’s corrected t-test p-value of 0.47). In the ‘novel chemistry’ case, the model had on average a balanced accuracy of 0.65, below the 31.6µM morphological profile classifier’s average balanced accuracy of 0.72.

To understand why in the ‘novel chemistry’ case, this model did not yield better performances, we computed the mean Morgan fingerprint Tanimoto distances between each chemical compound of the training set and its nearest neighbor in the training set, and the mean distances between each chemical compound of the testing set and its nearest neighbor in the training set.

In the ‘known chemistry’ case, on average, in the training set, each compound has a distance to its nearest neighbor of 0.50 and in the testing set 0.48. For the ‘novel chemistry’ case, on average, in the training set, each compound had a distance to its nearest neighbor of 0.49, and in the testing set 0.73.

In the ‘novel chemistry’ case, the training set did not have enough examples of distant chemical structures. To help the model, we added synthetic examples of distant chemical structures in the training set. To do so, we subset in each training set the cases where the predictions of the chemical structure similarity-based did not match the real class, and we updated the distances of the nearest neighbors with a random number between 0.7 and 0.9 and added those as synthetic examples in the dataset used to train the model.

By applying this approach, in the ‘known chemistry’ case the model achieved an average balanced accuracy of 0.85, slightly above, the chemical structure similarity-based classifier’s average balanced accuracy of 0.82 but was not significantly different (Nadeau and Bengio’s corrected t-test p-value of 0.12). In the ‘novel chemistry’ case, the model achieved an average balanced accuracy of 0.72 (Table 3).

## Discussion

Our results showed that the classification of compounds as very acute oral toxic or not, using a similarity-based approach, was possible using chemical structure information, morphological profiles of U2OS cells or the combination of both. When classifying compounds coming from the same chemical space as those in the classifier’s training set, the chemical structure information was more predictive. Conversely, when the compounds to be classified came from a different chemical space than those in the classifier’s training set, the morphological profiles of the U2OS cells were more predictive.

Initial attempts to use the publicly available QSAR model, CATMoS, for the prediction of acute oral toxicity on a set of 630 Bayer compounds did not yield good predictions ^5^. The CATMoS performance is hindered as Bayer Crop Science chemistry could be considered to be locally outside its training set (Table 2b). Although almost all compounds were globally within the CATMoS applicability domain, most resided in gaps of the training chemical space. It is well known that QSAR models excel when the compounds to be classified fall within their applicability domain of the models, and can perform poorly when they do not ^34^. In summary, CATMoS, which is a QSAR model trained on more than 10,000 compounds, has very good performance for the prediction of acute oral toxicity of those chemicals, but it does not work as effectively with the structurally diverse BCS chemistry.

To test our hypothesis, we trained a simple KNN classifier, resembling a similarity-based (or read-across) approach, on this set of 630 BCS compounds, using their chemical structure information. Working with two data-holdout strategies to simulate scenarios within and outside the applicability domain of a QSAR model, we evaluated our classifiers under two conditions: the ‘known chemistry’ case, simulating scenarios within applicability domain case, and the ‘novel chemistry’ case, attempting to simulate outside applicability domain case. In the ‘known chemistry’ case, our classifier exhibited strong performance, comparable to CATMoS :CATMoS achieved a balanced accuracy of 0. 84, in classifying compounds as very toxic (VT) (LD50 < 50 mg/kg) whereas our classifier had a balanced accuracy of 0.81 (Table 2c) for the classification of compounds as VAOT (LD50 < 60 mg/kg) ^5^.

However, as designed, the performance of the classifier dropped in the ‘novel chemistry’ case due to the data-holdout strategy, which placed Butina compound clusters, which are not present in the training sets, into the test sets. This effectively simulated scenarios outside of the models applicability domain, although the decrease in balanced accuracy (from 0.82 to 0.60) (Table 3) was not as drastic as that observed with the Bayer CropScience chemistry using CATMoS (from 0.84 to 0.52) (Table 2a).

To overcome this chemical applicability domain limitation, we explored whether using the compound biological effects could mitigate this problem. Compound-induced biological effects characterized by transcriptomics have previously been used to predict target activities, in association and comparison with QSAR models^35,36^. Here we utilized Cell Painting to generate morphological profiles at a more reasonable cost compared to transcriptomics.

Using a smaller set of 226 compounds, (balanced with 49 % of compounds where LD50<60 mg/kg; 51% where LD50>60 mg/kg), we trained KNN classifiers based on either chemical structure information or U2OS morphological profiles at 3 concentrations (10, 31.6 or 100 µM).

Morphological profiles at 31.6 µM concentration demonstrated better performance, compared to the other concentrations, in both the ‘known chemistry’ and ‘novel chemistry’ cases. With a balanced accuracy of 0.74 (Table 3) in the ‘known chemistry’ case and 0.72 (Table 3) in the ‘novel chemistry’ case, Cell Painting U2OS profiles demonstrated the ability to predict acute oral toxicity classes, interestingly, independent of the structural similarity of the tested compounds.

Cell Painting can indeed identify morphological patterns associated with specific mode of action (MoA) and molecular initiation event (MIE) of compounds ^37,38^. Typically, acute toxicity involves a limited number of MIEs ^39^. such as narcosis (activity at the lipid bilayer of the membrane), acetylcholinesterase inhibition, ion channel modulators and inhibitors of cellular respiration ^40^.

The Cell Painting experiment revealed morphological profiles (initiated by MIE) associated with acute oral toxicity, as evidenced by the grouping of morphological profiles associated with VAOT compounds. For example, four carbamates (Promecarb, Methiocarb, Propoxur, m- Cumenyl methylcarbamate) known as acetylcholinesterase inhibitors, produced similar morphological profiles in U2OS cells (Figure 8, middle row). This partially explains why the morphological profile-based classifiers were able to correctly classify the compounds. In the ‘known chemistry’ case, the performance of the classifiers based on the 31.6 µM morphological profile did not surpass the classifiers based on the chemical structure information but outperformed them in the ‘novel chemistry’ case.

The comparison of the two chemical structure based KNNs for the two datasets showed similar performances, but this comparison is limited because the two datasets are different. The QSAR dataset consisting of 630 compounds is unbalanced (109 VAOT, 521 NVAOT), while the Cell Painting dataset, made of 226 compounds is balanced, but of smaller size.

Capturing the biological effects of compounds had limitations: the limitation of the cell system to reveal the effects causally related to acute toxicity, together with technical limitations of the laboratory experiment itself.

For the limitation of the cell system, we observed, by Grit score analysis, that not all the compounds induced a biological response in U2OS cells (10% of the tested compounds), regardless of the concentration used. Six known VAOT compounds did not elicit any morphological changes compared to the negative controls. Among these compounds, five were public compounds and some of them with information on their possible mechanism of action. The warfarin, a vitamin K antagonist, and the methamidophos, a potent acetylcholinesterase inhibitor, did not induce any biological response in U2OS cells. This suggests that U2OS cells have their own biological applicability domain and may not capture all the bioactivities associated with acute oral toxicity observed in a whole organism such as a rat in our case study. Nevertheless, for our set of compounds, Cell Painting on U2OS managed to capture bioactivities for most of the VAOT compounds.

On the contrary, when analyzing the number of cells, we could also identify a limitation due to the cytotoxicity of the compounds: 44 compounds showed cytotoxicity at least at one concentration, and 12 exhibited cytotoxicity even at the lowest concentration. The morphological profiles of the cytotoxic treatments were not informative as they consisted mainly of debris and dying cells. It appears that the 31.6 µM concentration represented a good compromise between inducing bioactivity and avoiding cytotoxicity. However, building a model using only this concentration is a limitation, as morphological profiles coming from other concentrations could also have been associated with acute oral toxicity. Different rules for selecting the best concentration per compound, using the Grit score to ensure that the compound was active, or the number of cells to ensure that the compound was not cytotoxic did not produce better models.

For the limitation of the experiments, several quality issues can arise when conducting an experiment in a laboratory. Experiments are technically demanding and prone to variability and error. Seeding variability can affect the cell morphologies and thus morphological profiles.

There are other common problems that can occur in laboratory experiments, such as treatment errors, compounds with low purity or precipitation at high concentrations. These problems can affect the quality of the morphological profiles and thus the performance of a classifier based on morphological profile similarities.

The chemical structural information did not suffer from these limitations, because this information was not subject to quality issues, was not cell system dependent, and was not assay design dependent. This information was intrinsic to the description of a given compound. This may partly explain why chemical structure similarity-based classifiers, in the ‘known chemistry’ case, performed better than biological based classifiers: the full structural information is available, whereas the biological information is partially available and subject to quality issues, in particular reproducibility.

For both types of input data, the optimal number of neighbors for the KNN algorithm (supplementary information, S4) was 1, indicating that few examples of identified profiles leading to high acute oral toxicity or not were present in the dataset. A larger set of compounds, such as the CATMoS training set, would help to identify more examples of cell painting profiles associated with acute oral toxicity.

In the ‘novel chemistry’ case, the morphological profile-based classifiers did not experience as large a performance drop as the chemical structure similarity-based classifiers, suggesting that the biological space did not cluster in the same way as the chemical space. This indicates that similar compounds did not consistently induce the same response in U2OS cells (for example due to activity cliffs), and vice versa. The presence of different Butina clusters in the training and test sets did not necessarily result in different morphological profiles, explaining why the morphological profile-based classifier performance did not drop drastically in the ‘novel chemistry’ case.

The use of biological responses of compounds could be also an advantage with respect to enantiomers. The Morgan fingerprint used in this analysis does not take chirality into account. Enantiomers may have different acute oral toxicity, and the classifier based on chemical structure will not distinguish between them, where morphological profiles may be different.

The decision support model combined both predictions along with the nearest neighbor distances to make the final predictions, slightly improving the classification performance in the ‘known chemistry’ case but decreasing in the ‘novel chemistry’ case. By adding a few synthetic examples in each training set with higher distances in the chemical spaces, it was possible to increase the classification accuracy in the ‘known chemistry’ case, but not in the ‘novel chemistry’ case, where the model performed like the 31.6µM morphological-based classifier. Notably, in the ‘novel chemistry’ case the classifier preferred the predictions of the 31.6µM morphological-based classifier predictions over the predictions of the chemical structure similarity-based classifier.

Further results could extend and refine these findings by employing a broader set of compounds covering additional molecular initiating events (MIE) associated with acute oral toxicity. Additionally, a larger set of compounds could facilitate the identification of additional morphological profiles associated with acute oral toxicity. A larger dataset would also allow isolation of a set of compounds as an external dataset to further evaluate the performance of the classifier.

The choice of the KNN algorithm in this analysis was deliberate due to its simplicity and similarity to read-across approach commonly used in toxicology. Since the amount of in vivo data is often limited, the read-across approach is often the only analysis that can be performed. For chemical structure-based classifiers, other algorithms yielded similar performances (Supplementary Information).

To support the creation of public QSAR models with a wider applicability domain, representations of compound structures and results of acute toxicity studies for early candidates which failed to be placed on the markets could be shared by companies and organizations to expand the chemical space coverage.

In addition, in this analysis, the Morgan fingerprint was the only computed chemical fingerprint. The use of additional fingerprint or descriptors could help to achieve better QSAR and chemical structure similarity-based classifier performance.

Similarly, hand-crafted morphological features were used in this analysis. To capture a broader representation of morphological profiles, other deep learning based representations could also be tested^41^.

The decision support model uses the nearest neighbor distance to decide on which prediction to select. Other metrics, such as the ‘distance to model’, which is used to estimate a prediction uncertainty could also be used to decide on which prediction to use^42^.

We have also seen the limitation of the U2OS cell line with not capturing all the bioactivities of the compounds. Trying different cell lines could allow capturing more bioactivities linked to MIE leading to acute oral toxicity. Several cell lines have already been used with Cell Painting ^43,44^ and could help define a set of cell lines capable of capturing a maximum, if not all, MIE leading to acute oral toxicity.

Finally, absorption, distribution, metabolism, and excretion (ADME) properties of compounds were not taken into consideration in this study but incorporating such data could enhance predictive models. We tried to use predicted maximum concentration in plasma and AUC from a predictive model ^45^, but this information did not improve our results (data not shown).

In addition, pre-incubation of the compounds with liver S9 fractions (the 9000g supernatant of a liver homogenate), containing phase I and II metabolic enzymes, to generate the possible metabolites of a parent compound, could be helpful when the toxicity is driven by a metabolite, as it is done for example in the Ames assay to test the mutagenic potential of chemical compounds ^46,47^.

In conclusion, a combined approach utilizing chemical structure, and Cell Painting morphological profiles-based classifiers based on chemical and biological spaces distances holds promise for predicting acute oral toxicity. These classifiers could be used in the context of early de-risking and in the future serve in the context of Next Generation Risk Assessment (NGRA), which aims at refining if not replacing laboratory animal testing.

## Acknowledgment

The authors thank Dr. Nina Hallmark and Dr. Oscar Mendes Lucio for their insightful suggestions to improve the article.

## Data Availability

GitHub (https://github.com/Bayer-Group/cellPainting_acuteTox)

Consensus Cell Painting morphological profiles with oral acute toxicity label, and code are provided.

## Supporting Information

Lisf of reference compounds, list of public compounds, Umap of the chemical space with Butina clusters, other classifiers results, detailed CATMoS predictions, and python package versions.

## Abbreviation table

AD: Applicability Domain
BA: Balanced Accuracy
BCS: Bayer Crop Science
DMSO: Dimethyl sulfoxide
DS: Decision Support
GHS: Global Harmonized System
KNN: K Nearest Neighbor
LD50: Median Lethal Dose
MAD: Median Absolute Deviation
MCC: Matthews Correlation Coefficient
MIE: Molecular Initiative Event
MoA: Mode of Action
NGRA: Next Generation Risk Assessment
NVAOT: Non Very Acutely Oral Toxic (LD50 > 60 mg/kg)
QSAR: quantitative structure activity relationship
SMILES: Simplified molecular-input line-entry system
SN: Sensitivity
SP: Specificity
TP, TN, FP, FN: True Positives, True Negatives, False Positives, False Negatives
VAOT: Very Acutely Oral Toxic (LD50 ≤ 60 mg/kg)

## References

(1) Khalak, Y.; Tresadern, G.; Hahn, D. F.; De Groot, B. L.; Gapsys, V. Chemical Space Exploration with Active Learning and Alchemical Free Energies. J. Chem. Theory Comput. 2022, 18 (10), 6259–6270. 10.1021/acs.jctc.2c00752.

(2) Scannell, J. W.; Bosley, J.; Hickman, J. A.; Dawson, G. R.; Truebel, H.; Ferreira, G. S.; Richards, D.; Treherne, J. M. Predictive Validity in Drug Discovery: What It Is, Why It Matters and How to Improve It. Nat Rev Drug Discov 2022. 10.1038/s41573-022-00552-x.

(3) Henriquez, J. E.; Badwaik, V. D.; Bianchi, E.; Chen, W.; Corvaro, M.; LaRocca, J.; Lunsman, T. D.; Zu, C.; Johnson, K. J. From Pipeline to Plant Protection Products: Using New Approach Methodologies (NAMs) in Agrochemical Safety Assessment. J. Agric. Food Chem. 2024, 72 (19), 10710–10724. 10.1021/acs.jafc.4c00958.

(4) Erhirhie, E. O.; Ihekwereme, C. P.; Ilodigwe, E. E. Advances in Acute Toxicity Testing: Strengths, Weaknesses and Regulatory Acceptance. Interdisciplinary Toxicology 2018, 11 (1), 5–12. 10.2478/intox-2018-0001.

(5) Mansouri, K.; Karmaus, A. L.; Fitzpatrick, J.; Patlewicz, G.; Pradeep, P.; Alberga, D.; Alepee, N.; Allen, T. E. H.; Allen, D.; Alves, V. M.; Andrade, C. H.; Auernhammer, T. R.; Ballabio, D.; Bell, S.; Benfenati, E.; Bhattacharya, S.; Bastos, J. V.; Boyd, S.; Brown, J. B.; Capuzzi, S. J.; Chushak, Y.; Ciallella, H.; Clark, A. M.; Consonni, V.; Daga, P. R.; Ekins, S.; Farag, S.; Fedorov, M.; Fourches, D.; Gadaleta, D.; Gao, F.; Gearhart, J. M.; Goh, G.; Goodman, J. M.; Grisoni, F.; Grulke, C. M.; Hartung, T.; Hirn, M.; Karpov, P.; Korotcov, A.; Lavado, G. J.; Lawless, M.; Li, X.; Luechtefeld, T.; Lunghini, F.; Mangiatordi, G. F.; Marcou, G.; Marsh, D.; Martin, T.; Mauri, A.; Muratov, E. N.; Myatt, G. J.; Nguyen, D.-T.; Nicolotti, O.; Note, R.; Pande, P.; Parks, A. K.; Peryea, T.; Polash, A. H.; Rallo, R.; Roncaglioni, A.; Rowlands, C.; Ruiz, P.; Russo, D. P.; Sayed, A.; Sayre, R.; Sheils, T.; Siegel, C.; Silva, A. C.; Simeonov, A.; Sosnin, S.; Southall, N.; Strickland, J.; Tang, Y.; Teppen, B.; Tetko, I. V.; Thomas, D.; Tkachenko, V.; Todeschini, R.; Toma, C.; Tripodi, I.; Trisciuzzi, D.; Tropsha, A.; Varnek, A.; Vukovic, K.; Wang, Z.; Wang, L.; Waters, K. M.; Wedlake, A. J.; Wijeyesakere, S. J.; Wilson, D.; Xiao, Z.; Yang, H.; Zahoranszky-Kohalmi, G.; Zakharov, A. V.; Zhang, F. F.; Zhang, Z.; Zhao, T.; Zhu, H.; Zorn, K. M.; Casey, W.; Kleinstreuer, N. C. CATMoS: Collaborative Acute Toxicity Modeling Suite. Environ Health Perspect 2021, 129 (4), 047013. 10.1289/EHP8495.

(6) Bray, M.-A.; Singh, S.; Han, H.; Davis, C. T.; Borgeson, B.; Hartland, C.; Kost-Alimova, M.; Gustafsdottir, S. M.; Gibson, C. C.; Carpenter, A. E. Cell Painting, a High-Content Image- Based Assay for Morphological Profiling Using Multiplexed Fluorescent Dyes. Nat Protoc 2016, 11 (9), 1757–1774. 10.1038/nprot.2016.105.

(7) Chandrasekaran, S. N.; Ceulemans, H.; Boyd, J. D.; Carpenter, A. E. Image-Based Profiling for Drug Discovery: Due for a Machine-Learning Upgrade? Nat Rev Drug Discov 2021, 20 (2), 145–159. 10.1038/s41573-020-00117-w.

(8) Lapins, M.; Spjuth, O. Evaluation of Gene Expression and Phenotypic Profiling Data as Quantitative Descriptors for Predicting Drug Targets and Mechanisms of Action. March 17, 2019. 10.1101/580654.

(9) Tian, G.; Harrison, P. J.; Sreenivasan, A. P.; Carreras-Puigvert, J.; Spjuth, O. Combining Molecular and Cell Painting Image Data for Mechanism of Action Prediction. Artificial Intelligence in the Life Sciences 2023, 3, 100060. 10.1016/j.ailsci.2023.100060.

(10) Simm, J.; Klambauer, G.; Arany, A.; Steijaert, M.; Wegner, J. K.; Gustin, E.; Chupakhin, V.; Chong, Y. T.; Vialard, J.; Buijnsters, P.; Velter, I.; Vapirev, A.; Singh, S.; Carpenter, A. E.; Wuyts, R.; Hochreiter, S.; Moreau, Y.; Ceulemans, H. Repurposing High-Throughput Image Assays Enables Biological Activity Prediction for Drug Discovery. Cell Chemical Biology 2018, 25 (5), 611–618.e3. 10.1016/j.chembiol.2018.01.015.

(11) Nyffeler, J.; Willis, C.; Lougee, R.; Richard, A.; Paul-Friedman, K.; Harrill, J. A. Bioactivity Screening of Environmental Chemicals Using Imaging-Based High-Throughput Phenotypic Profiling. Toxicology and Applied Pharmacology 2020, 389, 114876. 10.1016/j.taap.2019.114876.

(12) Garcia De Lomana, M.; Marin Zapata, P. A.; Montanari, F. Predicting the Mitochondrial Toxicity of Small Molecules: Insights from Mechanistic Assays and Cell Painting Data. Chem. Res. Toxicol. 2023, 36 (7), 1107–1120. 10.1021/acs.chemrestox.3c00086.

(13) Seal, S.; Carreras-Puigvert, J.; Trapotsi, M.-A.; Yang, H.; Spjuth, O.; Bender, A. Integrating Cell Morphology with Gene Expression and Chemical Structure to Aid Mitochondrial Toxicity Detection. Commun Biol 2022, 5 (1), 858. 10.1038/s42003-022-03763-5.

(14) Lejal, V.; Cerisier, N.; Rouquié, D.; Taboureau, O. Assessment of Drug-Induced Liver Injury through Cell Morphology and Gene Expression Analysis. Chem. Res. Toxicol. 2023, 36 (9), 1456–1470. 10.1021/acs.chemrestox.2c00381.

(15) Kowalski, B. R.; Bender, C. F. K-Nearest Neighbor Classification Rule (Pattern Recognition) Applied to Nuclear Magnetic Resonance Spectral Interpretation. Anal. Chem. 1972, 44 (8), 1405–1411. 10.1021/ac60316a008.

(16) Zhu, H. Supporting Read-across Using Biological Data. ALTEX 2016, 167–182. 10.14573/altex.1601252.

(17) ChemIDplus, 2023. https://pubchem.ncbi.nlm.nih.gov/source/ChemIDplus.

(18) Butina, D. Unsupervised Data Base Clustering Based on Daylight’s Fingerprint and Tanimoto Similarity: A Fast and Automated Way To Cluster Small and Large Data Sets. J. Chem. Inf. Comput. Sci. 1999, 39 (4), 747–750. 10.1021/ci9803381.

(19) Mansouri, K.; Grulke, C. M.; Judson, R. S.; Williams, A. J. OPERA Models for Predicting Physicochemical Properties and Environmental Fate Endpoints. J Cheminform 2018, 10 (1), 10. 10.1186/s13321-018-0263-1.

(20) Cimini, B. A.; Chandrasekaran, S. N.; Kost-Alimova, M.; Miller, L.; Goodale, A.; Fritchman, B.; Byrne, P.; Garg, S.; Jamali, N.; Logan, D. J.; Concannon, J. B.; Lardeau, C.-H.; Mouchet, E.; Singh, S.; Shafqat Abbasi, H.; Aspesi, P.; Boyd, J. D.; Gilbert, T.; Gnutt, D.; Hariharan, S.; Hernandez, D.; Hormel, G.; Juhani, K.; Melanson, M.; Mervin, L. H.; Monteverde, T.; Pilling, J. E.; Skepner, A.; Swalley, S. E.; Vrcic, A.; Weisbart, E.; Williams, G.; Yu, S.; Zapiec, B.; Carpenter, A. E. Optimizing the Cell Painting Assay for Image-Based Profiling. Nat Protoc 2023, 18 (7), 1981–2013. 10.1038/s41596-023-00840-9.

(21) Stirling, D. R.; Swain-Bowden, M. J.; Lucas, A. M.; Carpenter, A. E.; Cimini, B. A.; Goodman, A. CellProfiler 4: Improvements in Speed, Utility and Usability. BMC Bioinformatics 2021, 22 (1), 433. 10.1186/s12859-021-04344-9.

(22) Serrano, E.; Chandrasekaran, S. N.; Bunten, D.; Brewer, K. I.; Tomkinson, J.; Kern, R.; Bornholdt, M.; Fleming, S.; Pei, R.; Arevalo, J.; Tsang, H.; Rubinetti, V.; Tromans-Coia, C.; Becker, T.; Weisbart, E.; Bunne, C.; Kalinin, A. A.; Senft, R.; Taylor, S. J.; Jamali, N.; Adeboye, A.; Abbasi, H. S.; Goodman, A.; Caicedo, J. C.; Carpenter, A. E.; Cimini, B. A.; Singh, S.; Way, G. P. Reproducible Image-Based Profiling with Pycytominer. 2023. 10.48550/ARXIV.2311.13417.

(23) Benchmarking Grit. Benchmarking Grit. https://github.com/broadinstitute/grit-benchmark (accessed 2024-05-29).

(24) Cytominer-eval: Evaluating quality of perturbation profiles. Cytominer-eval: Evaluating quality of perturbation profiles. https://github.com/cytomining/cytominereval (accessed 2024-05-29).

(25) Morgan, H. L. The Generation of a Unique Machine Description for Chemical Structures-A Technique Developed at Chemical Abstracts Service. J. Chem. Doc. 1965, 5 (2), 107–113. 10.1021/c160017a018.

(26) Rogers, D.; Hahn, M. Extended-Connectivity Fingerprints. J. Chem. Inf. Model. 2010, 50 (5), 742–754. 10.1021/ci100050t.

(27) Landrum, G.; Tosco, P.; Kelley, B.; Ric; Cosgrove, D.; Sriniker; Gedeck; Vianello, R.; NadineSchneider; Kawashima, E.; N, D.; Jones, G.; Dalke, A.; Cole, B.; Swain, M.; Turk, S.; AlexanderSavelyev; Vaucher, A.; Wójcikowski, M.; Ichiru Take; Probst, D.; Ujihara, K.; Scalfani, V. F.; Godin, G.; Lehtivarjo, J.; Walker, R.; Pahl, A.; Francois Berenger; Jasondbiggs; Strets123. Rdkit/Rdkit: 2023_03_3 (Q1 2023) Release, 2023. 10.5281/ZENODO.591637.

(28) Pedregosa, F.; Varoquaux, G.; Gramfort, A.; Michel, V.; Thirion, B.; Grisel, O.; Blondel, M.; Prettenhofer, P.; Weiss, R.; Dubourg, V.; Vanderplas, J.; Passos, A.; Cournapeau, D.; Brucher, M.; Perrot, M.; Duchesnay, E. Scikit-Learn: Machine Learning in Python. Journal of Machine Learning Research 2011, 12, 2825–2830.

(29) Escher, S. E.; Bitsch, A. Read-Across Methodology in Toxicological Risk Assessment. In Regulatory Toxicology; Reichl, F.-X., Schwenk, M., Eds.; Springer Berlin Heidelberg: Berlin, Heidelberg, 2021; pp 1–14. 10.1007/978-3-642-36206-4_132-1.

(30) Seal, S.; Yang, H.; Trapotsi, M.-A.; Singh, S.; Carreras-Puigvert, J.; Spjuth, O.; Bender, A. Merging Bioactivity Predictions from Cell Morphology and Chemical Fingerprint Models Using Similarity to Training Data. J Cheminform 2023, 15 (1), 56. 10.1186/s13321-023-00723-x.

(31) Cristianini, N.; Ricci, E. Support Vector Machines: 1992; Boser, Guyon, Vapnik. In Encyclopedia of Algorithms; Kao, M.-Y., Ed.; Springer US: Boston, MA, 2008; pp 928–932. 10.1007/978-0-387-30162-4_415.

(32) Nadeau, C.; Bengio, Y. Inference for the Generalization Error. Machine Learning 2003, 52 (3), 239–281. 10.1023/A:1024068626366.

(33) Sainburg, T.; McInnes, L.; Gentner, T. Q. Parametric UMAP Embeddings for Representation and Semisupervised Learning. Neural Computation 2021, 33 (11), 2881–2907.

(34) Kar, S.; Roy, K.; Leszczynski, J. Applicability Domain: A Step Toward Confident Predictions and Decidability for QSAR Modeling. In Computational Toxicology; Nicolotti, O., Ed.; Methods in Molecular Biology; Springer New York: New York, NY, 2018; Vol. 1800, pp 141–169. 10.1007/978-1-4939-7899-1_6.

(35) Baillif, B.; Wichard, J.; Méndez-Lucio, O.; Rouquié, D. Exploring the Use of Compound- Induced Transcriptomic Data Generated From Cell Lines to Predict Compound Activity Toward Molecular Targets. Front. Chem. 2020, 8, 296. 10.3389/fchem.2020.00296.

(36) Moshkov, N.; Becker, T.; Yang, K.; Horvath, P.; Dancik, V.; Wagner, B. K.; Clemons, P. A.; Singh, S.; Carpenter, A. E.; Caicedo, J. C. Predicting Compound Activity from Phenotypic Profiles and Chemical Structures. Nat Commun 2023, 14 (1), 1967. 10.1038/s41467-023-37570-1.

(37) Ljosa, V.; Caie, P. D.; Ter Horst, R.; Sokolnicki, K. L.; Jenkins, E. L.; Daya, S.; Roberts, M. E.; Jones, T. R.; Singh, S.; Genovesio, A.; Clemons, P. A.; Carragher, N. O.; Carpenter, A. E. Comparison of Methods for Image-Based Profiling of Cellular Morphological Responses to Small-Molecule Treatment. J Biomol Screen 2013, 18 (10), 1321–1329. 10.1177/1087057113503553.

(38) Way, G. P.; Natoli, T.; Adeboye, A.; Litichevskiy, L.; Yang, A.; Lu, X.; Caicedo, J. C.; Cimini, B. A.; Karhohs, K.; Logan, D. J.; Rohban, M. H.; Kost-Alimova, M.; Hartland, K.; Bornholdt, M.; Chandrasekaran, S. N.; Haghighi, M.; Weisbart, E.; Singh, S.; Subramanian, A.; Carpenter, A. E. Morphology and Gene Expression Profiling Provide Complementary Information for Mapping Cell State. Cell Systems 2022, 13 (11), 911–923.e9. 10.1016/j.cels.2022.10.001.

(39) Prieto, P. Investigating Cell Type Specific Mechanisms Contributing to Acute Oral Toxicity. ALTEX 2019, 36 (1), 39–64. 10.14573/altex.1805181.

(40) A. Leblanc, G. Mechanisms of Acute Toxicity. In A Textbook of Modern Toxicology; 2004.

(41) Caicedo, J. C.; McQuin, C.; Goodman, A.; Singh, S.; Carpenter, A. E. Weakly Supervised Learning of Single-Cell Feature Embeddings. In 2018 IEEE/CVF Conference on Computer Vision and Pattern Recognition; IEEE: Salt Lake City, UT, 2018; pp 9309–9318. 10.1109/CVPR.2018.00970.

(42) Sushko, I.; Novotarskyi, S.; Körner, R.; Pandey, A. K.; Cherkasov, A.; Li, J.; Gramatica, P.; Hansen, K.; Schroeter, T.; Müller, K.-R.; Xi, L.; Liu, H.; Yao, X.; Öberg, T.; Hormozdiari, F.; Dao, P.; Sahinalp, C.; Todeschini, R.; Polishchuk, P.; Artemenko, A.; Kuz’min, V.; Martin, T. M.; Young, D. M.; Fourches, D.; Muratov, E.; Tropsha, A.; Baskin, I.; Horvath, D.; Marcou, G.; Muller, C.; Varnek, A.; Prokopenko, V. V.; Tetko, I. V. Applicability Domains for Classification Problems: Benchmarking of Distance to Models for Ames Mutagenicity Set. J. Chem. Inf. Model. 2010, 50 (12), 2094–2111. 10.1021/ci100253r.

(43) Cox, M. J.; Jaensch, S.; Van de Waeter, J.; Cougnaud, L.; Seynaeve, D.; Benalla, S.; Koo, S. J.; Van Den Wyngaert, I.; Neefs, J.-M.; Malkov, D.; Bittremieux, M.; Steemans, M.; Peeters, P. J.; Wegner, J. K.; Ceulemans, H.; Gustin, E.; Chong, Y. T.; Göhlmann, H. W. H. Tales of 1,008 Small Molecules: Phenomic Profiling through Live-Cell Imaging in a Panel of Reporter Cell Lines. Sci Rep 2020, 10 (1), 13262. 10.1038/s41598-020-69354-8.

(44) Nyffeler, J. Phenotypic Profiling for High-Throughput Chemical Bioactivity Screening at the U.S. EPA. 2020, 11457644 Bytes. 10.23645/EPACOMPTOX.13198346.

(45) Schneckener, S.; Grimbs, S.; Hey, J.; Menz, S.; Osmers, M.; Schaper, S.; Hillisch, A.; Göller, A. H. Prediction of Oral Bioavailability in Rats: Transferring Insights from in Vitro Correlations to (Deep) Machine Learning Models Using in Silico Model Outputs and Chemical Structure Parameters. J. Chem. Inf. Model. 2019, 59 (11), 4893–4905. 10.1021/acs.jcim.9b00460.

(46) Hakura, A.; Suzuki, S.; Sawada, S.; Motooka, S.; Satoh, T. An Improvement of the Ames Test Using a Modified Human Liver S9 Preparation. Journal of Pharmacological and Toxicological Methods 2001, 46 (3), 169–172. 10.1016/S1056-8719(02)00186-7.

(47) Hopperstad, K.; DeGroot, D. E.; Zurlinden, T.; Brinkman, C.; Thomas, R. S.; Deisenroth, C. Chemical Screening in an Estrogen Receptor Transactivation Assay With Metabolic Competence. Toxicological Sciences 2022, 187 (1), 112–126. 10.1093/toxsci/kfac019.

