## Supplementary information for "Cell Painting and chemical structure read-across can complement each other for rat acute oral toxicity prediction in chemical early de-risking" (from another version of this preprint: version 1, 2024-04-28)

| Chemical name | Concentration (μM) |
|---|---|
| staurosporine | 0.0316 |
| staurosporine | 0.01 |
| staurosporine | 0.1 |
| staurosporine | 0.316 |
| staurosporine | 1.0 |
| TCS7004 | 10.0 |
| rotenone | 1.0 |
| rosiglitazone | 100.0 |
| NVS-PAK1 | 10.0 |
| LY2109761 | 10.0 |
| FK886 | 10.0 |
| dexamethasone | 10.0 |
| berberine | 10.0 |
| AMG-900 | 10.0 |

**Table S1**.

Positive reference compounds

| CASRN | Chemical Name | Rat acute oral LD50 | Class |
|---|---|---|---|
| 2631-37-0 | Promecarb | 35.0 | Insecticide (carbamate) |
| 6610-29-3 | N-Methylhydrazinecarbothioamide | 14.0 | |
| 2032-65-7 | Methiocarb | 20.0 | Insecticide (carbamate) |
| 2231-57-4 | Thiocarbazide | 21.6 | USED IN ELECTRON MICROSCOPY |
| 931-97-5 | Cyclohexanone cyanohydrin | 10.0 | NA |
| 786-19-6 | Carbophenothion | 6.8 | Insecticide |
| 53-86-1 | Indomethacin | 2.42 | A non-steroidal anti-inflammatory drug |
| 53164-05-9 | Acemetacin | 24.2 | A non-steroidal anti-inflammatory drug |
| 114-26-1 | Propoxur | 41.0 | Insecticide (carbamate) |
| 83-26-1 | Pindone | 10.3 | Rodenticide. Anticoagulant |
| 1420-07-1 | Dinoterb | 26.0 | Herbicide and rodenticide |
| 103-85-5 | 1-Phenyl-2-thiourea | 3.0 | Used to make rodenticide (thiourea) |
| 10265-92-6 | Methamidophos | 9.25 | Insecticide |
| 2610-86-8 | Potassium warfarin | 58.0 | Rodenticide. Anticoagulant. inhibiting the synthesis of vitamin K-dependent coagulation factors |
| 1744-22-5 | Riluzole | 45.0 | Anticonvulsant. Glutamate antagonist. |
| 107-49-3 | Tetraethyl pyrophosphate | 0.5 | Pesticide. inhibitor of cholinesterase |
| 3333-52-6 | Tetramethylsuccinonitrile | 27.0 | (decomposition product) |
| 541-53-7 | 2,4-Dithiobiuret | 5.0 | plasticizer, rubber accelerator, and resin intermediate; Also used to make insecticides and rodenticides and to delay wilting of flowers |
| 87-86-5 | Pentachlorophenol | 38.5 | Pesticide. Fungicide. |
| 145-73-3 | Endothal | 38.0 | Pesticide. Herbicide. |
| 66-75-1 | Uracil mustard | 3.55 | anti-cancer medicine. inhibits DNA synthesis |
| 53558-25-1 | Pyrinuron | 4.75 | Pesticide. Rodenticide |
| 64-00-6 | m-Cumenyl methylcarbamate | 16.0 | Pesticide. Insecticide (carbamate) |
| 56-72-4 | Coumaphos | 13.0 | Pesticide. Insecticide |
| 973-21-7 | Dinobuton | 59.0 | Pesticide. Insecticide. Fungicide. |
| 82-66-6 | Diphacinone | 1.5 | rodenticide and anticoagulant medication |
| 15307-79-6 | Diclofenac sodium | 53.0 | nonsteroidal anti-inflammatory drug |
| 106266-06-2 | Risperidone | 56.6 | second-generation antipsychotic. serotonergic antagonist, alpha-adrenergic antagonist, H1-receptor antagonist, dopaminergic antagonist, EC 3.4.21.26 (prolyl oligopeptidase) inhibitor. |
| 1491-59-4 | Oxymetazoline | 0.8 | Vasoconstrictor. alpha (α)-adrenergic agonist with affinity to both α1- and α2-adrenoceptors. |

**Table S2.**

Public chemical compounds with an acute oral LD50 below 60 mg/kg

**a: Classifiers using Morgan fingerprints**

| | Easy case | | | | | Difficult case | | | | |
|---|---|---|---|---|---|---|---|---|---|---|
| | **ACC** | **BA** | **MCC** | **SN** | **SP** | **ACC** | **BA** | **MCC** | **SN** | **SP** |
| **KNN** | 0.82 | 0.82 | 0.65 | 0.82 | 0.82 | 0.61 | 0.60 | 0.20 | 0.48 | 0.72 |
| **SVM** | 0.83 | 0.83 | 0.67 | 0.84 | 0.82 | 0.65 | 0.65 | 0.30 | 0.42 | 0.87 |
| **random forest** | 0.82 | 0.82 | 0.66 | 0.82 | 0.83 | 0.62 | 0.62 | 0.25 | 0.39 | 0.85 |
| **xg boost** | 0.78 | 0.78 | 0.56 | 0.79 | 0.77 | 0.57 | 0.57 | 0.13 | 0.39 | 0.76 |

**b: Classifiers using morphological profiles (31.6 µM)**

| | Easy case | | | | | Difficult case | | | | |
|---|---|---|---|---|---|---|---|---|---|---|
| | **ACC** | **BA** | **MCC** | **SN** | **SP** | **ACC** | **BA** | **MCC** | **SN** | **SP** |
| **KNN** | 0.75 | 0.75 | 0.51 | 0.71 | 0.79 | 0.70 | 0.72 | 0.42 | 0.68 | 0.76 |
| **SVM** | 0.69 | 0.69 | 0.40 | 0.58 | 0.80 | 0.63 | 0.63 | 0.26 | 0.47 | 0.79 |
| **random forest** | 0.66 | 0.66 | 0.33 | 0.62 | 0.70 | 0.58 | 0.59 | 0.18 | 0.51 | 0.67 |
| **xg boost** | 0.66 | 0.66 | 0.32 | 0.62 | 0.70 | 0.56 | 0.56 | 0.13 | 0.48 | 0.64 |

**Table S3.**

Mean over 100 splits of 5 metrics: Accuracy (ACC), Balanced Accuracy (BA), Matthew's correlation coefficient (MCC), Sensitivity (SN), Specificity ‘SP) for the classification of compounds as VAOT and NVAOT.4 classifiers were uses: K nearest neighbor (KNN), Support Vector Machine (SVM), Random Forest and xg Boost, for the easy and difficult cases.

a: classifier using Morgan fingerprints.

b: classifiers using morphological profiles of U2OS cells perturbed with chemical compounds at 31.6 µM

| | a) Classifiers using Morgan Fingerprints | | | | | | | | | | |
|---|---|---|---|---|---|---|---|---|---|---|---|
| | | Easy case | | | | | Difficult case | | | | |
| **weights** | **Number of neighboor** | **ACC** | **aBA** | **MCC** | **SP** | **SN** | **ACC** | **BA** | **MCC** | **SP** | **SN** |
| uniform | 1 | 0.82 | 0.82 | 0.65 | 0.82 | 0.82 | 0,62 | 0,62 | 0,22 | 0,50 | 0,73 |
| uniform | 3 | 0.79 | 0.79 | 0.60 | 0.79 | 0.80 | 0,61 | 0,60 | 0,21 | 0,44 | 0,77 |
| uniform | 5 | 0.77 | 0.77 | 0.55 | 0.77 | 0.78 | 0,60 | 0,61 | 0,20 | 0,48 | 0,74 |
| uniform | 7 | 0.73 | 0.73 | 0.47 | 0.74 | 0.73 | 0,57 | 0,57 | 0,14 | 0,41 | 0,73 |
| uniform | 9 | 0.72 | 0.72 | 0.46 | 0.71 | 0.74 | 0,56 | 0,57 | 0,13 | 0,39 | 0,75 |
| uniform | 11 | 0.71 | 0.71 | 0.42 | 0.69 | 0.73 | 0,56 | 0,56 | 0,11 | 0,38 | 0,74 |
| uniform | 13 | 0.70 | 0.70 | 0.40 | 0.65 | 0.74 | 0,54 | 0,54 | 0,08 | 0,35 | 0,74 |
| distance | 3 | 0.81 | 0.81 | 0.64 | 0.82 | 0.81 | 0,61 | 0,61 | 0,21 | 0,44 | 0,77 |
| distance | 5 | 0.81 | 0.81 | 0.64 | 0.83 | 0.80 | 0,60 | 0,61 | 0,20 | 0,48 | 0,74 |
| distance | 7 | 0.78 | 0.78 | 0.57 | 0.78 | 0.78 | 0,57 | 0,57 | 0,14 | 0,41 | 0,73 |
| distance | 9 | 0.77 | 0.77 | 0.54 | 0.75 | 0.78 | 0,56 | 0,57 | 0,13 | 0,39 | 0,75 |
| distance | 11 | 0.75 | 0.75 | 0.50 | 0.72 | 0.77 | 0,56 | 0,56 | 0,11 | 0,38 | 0,74 |
| distance | 13 | 0.72 | 0.72 | 0.45 | 0.69 | 0.76 | 0,54 | 0,54 | 0,08 | 0,35 | 0,74 |
| | b) KNN using morphological profiles (31.6 µM) | | | | | | | | | | |
| | | Easy case | | | | | Difficult case | | | | |

| weights | Number of neighboor | ACC | BA | MCC | SP | SN | ACC | BA | MCC | SP | SN |
|---|---|---|---|---|---|---|---|---|---|---|---|
| uniform | 1 | 0.75 | 0.75 | 0.51 | 0.71 | 0.79 | 0,69 | 0,72 | 0,42 | 0,68 | 0,76 |
| uniform | 3 | 0.68 | 0.68 | 0.37 | 0.65 | 0.72 | 0,64 | 0,64 | 0,27 | 0,58 | 0,71 |
| uniform | 5 | 0.68 | 0.68 | 0.37 | 0.64 | 0.72 | 0,62 | 0,63 | 0,25 | 0,59 | 0,68 |
| uniform | 7 | 0.68 | 0.68 | 0.37 | 0.65 | 0.71 | 0,60 | 0,62 | 0,22 | 0,56 | 0,67 |
| uniform | 9 | 0.67 | 0.67 | 0.34 | 0.63 | 0.70 | 0,57 | 0,59 | 0,17 | 0,53 | 0,65 |
| uniform | 11 | 0.64 | 0.64 | 0.29 | 0.62 | 0.66 | 0,56 | 0,58 | 0,14 | 0,53 | 0,62 |
| uniform | 13 | 0.65 | 0.65 | 0.30 | 0.62 | 0.67 | 0,60 | 0,61 | 0,20 | 0,57 | 0,65 |
| distance | 3 | 0.69 | 0.69 | 0.38 | 0.64 | 0.73 | 0,65 | 0,65 | 0,30 | 0,60 | 0,71 |
| distance | 5 | 0.68 | 0.68 | 0.36 | 0.63 | 0.72 | 0,63 | 0,64 | 0,26 | 0,59 | 0,68 |
| distance | 7 | 0.70 | 0.69 | 0.40 | 0.66 | 0.73 | 0,62 | 0,63 | 0,25 | 0,58 | 0,69 |
| distance | 9 | 0.69 | 0.69 | 0.38 | 0.64 | 0.73 | 0,60 | 0,61 | 0,22 | 0,55 | 0,68 |
| distance | 11 | 0.67 | 0.67 | 0.34 | 0.63 | 0.71 | 0,59 | 0,60 | 0,19 | 0,54 | 0,66 |
| distance | 13 | 0.68 | 0.68 | 0.36 | 0.63 | 0.72 | 0,63 | 0,64 | 0,26 | 0,59 | 0,69 |

**Table S4.**

Mean over 100 splits of 5 metrics: Accuracy (ACC), Balanced Accuracy (BA), Matthew's correlation coefficient (MCC), Sensitivity (SN), Specificity 'SP) for the classification of compounds as VAOT and NVAOT using KNN models, varying the number of neighbors and the weights, in the easy and difficult cases.

a) using Morgan fingerprint with Tanimoto distance.
b) using morphological profiles (31.6 µM) with correlation distance.

### Detailed CATMoS predictions

| | | | Predicted GHS categories | | | | |
|---|---|---|---|---|---|---|---|
| | | | I | II | III | IV | NC |
| | | **LD50** (mg/kg) | ≤5 | >5 ≤50 | >50 ≤300 | >300 ≤2000 | >2000 |
| **True categories** | **VOAT** | ≤ 60 | 0 | 5 | 15 | 29 | 60 |
| | **NVAOT** | >60 | 1 | 6 | 54 | 199 | 261 |

**Table S5.**

CATMoS GHS category predictions for the set of 630 Bayer compounds. The true categories are the categories VAOT (≤ 60 mg/kg) and NVAOT (> 60 mg/kg) of the Bayer compounds. The predicted GHS categories are I (≤ 5 mg/kg),, II (> 5 mg/kg ≤50 mg/kg),, III (>50 mg/kg ≤ 300 mg/kg), IV (>300 mg/kg≤2000 mg/kg) and NC (> 2000 mg/kg).
